# Structure of pre-miR-31 reveals an active role in Dicer processing

**DOI:** 10.1101/2023.01.03.519659

**Authors:** Sicong Ma, Anita Kotar, Scott Grote, Silvi Rouskin, Sarah C. Keane

## Abstract

As an essential post-transcriptional regulator of gene expression, microRNA (miR) levels must be strictly maintained. The biogenesis of many, but not all, miRs is mediated by trans-acting protein partners through a variety of mechanisms, including remodeling of the RNA structure. miR-31 functions as an oncogene in numerous cancers and interestingly, its biogenesis is not known to be regulated by protein binding partners. Therefore, the intrinsic structural properties of pre-miR-31 can provide a mechanism by which its biogenesis is regulated. We determined the solution structure of the precursor element of miR-31 (pre-miR-31) to investigate the role of distinct structural elements in regulating Dicer processing. We found that the presence or absence of mismatches within the helical stem do not strongly influence Dicer processing of the pre-miR. However, both the apical loop size and structure at the Dicing site are key elements for discrimination by Dicer. Interestingly, our NMR-derived structure reveals the presence of a triplet of base pairs that link the Dicer cleavage site and the apical loop. Mutational analysis in this region suggests that the stability of the junction region strongly influence both Dicer binding and processing. Our results enrich our understanding of the active role that RNA structure plays in regulating Dicer processing which has direct implications for control of gene expression.

## Introduction

MicroRNAs (miRs) are a class of small non-coding RNAs that regulate protein gene expression post-transcriptionally. By base pairing with target mRNAs, miRs trigger mRNA degradation or translational suppression^[1-4]^. Abnormal miRs levels are associated with cancers, diabetes, neurological and other diseases^[5-8]^. RNA polymerase II transcribes primary microRNA (pri-miR) in the nucleus and pri-miRs are subsequently processed by Microprocessor, which is composed of Drosha and DiGeorge syndrome critical region 8 (DGCR8) proteins, to generate precursor microRNAs (pre-miRs). Pre-miRs are exported from the nucleus to the cytoplasm in a GTP-dependent manner by Exportin-5. In the cytoplasm, pre-miRs are further processed by Dicer to generate 21-22 nucleotide (nt) mature miR duplexes^[4, 9]^. Argonaute (Ago) protein loads the miR duplex and subsequently displaces one of the strands from the complex to form the miR-induced silencing complex, which is responsible for mRNA degradation or translational suppression^[2, 4]^.

Distinctive regulatory elements for pri-miRs and pre-miRs have been discovered over past two decades. These elements include specific sequences within the pri-miRs and pre-miRs that recruit regulatory proteins^[10-13]^ and structural features of pri-miRs and pre-miRs that mediate enzymatic processing^[14-19]^. Although protein-mediated secondary structure^[10, 20]^ or primary sequence switches^[11]^ are largely correlated with the differential expression of mature miRs^[4]^, the specific secondary structure elements and/or structural plasticity of pri/pre-miR are both known to be intrinsic regulatory factors^[17, 19, 21]^. For example, Interleukin enhancer-binding factor 3 (ILF3) is a regulatory protein for pre-miR-144 dicing by reshaping the terminal loop to form a suboptimal substrate for Dicer processing^[20]^. Meanwhile, the Lin28 RNA binding protein (RBP) is a classic example of a protein which promotes pre-let-7 turnover by recruiting terminal uridyltransferase (TUTase) which promotes degradation of the pre-miR.^[12, 22]^ While protein-mediated regulation is indeed important for many pre-miRs, a recent study showed that pre-miR-21 exists as a pH-dependent two-state ensemble and excited pre-miR-21 is an efficient cleavage substrate for Dicer protein^[19, 23]^. Therefore, the intrinsic structural properties of a pre-miR may serve as an alternative mechanism for regulation of its biogenesis, suggesting that the RNA is not a passive element in miR biogenesis.

MicroRNA-31 (miR-31) acts as oncogene in multiple cancers. Upregulation of miR-31 in cells is associated with cancer proliferation, anti-apoptosis and migration in multiple cancers by targeting different biogenesis pathways in cells^[24]^. For example, in colorectal cancers (CRC), overexpression of miR-31 promotes cancer proliferation by targeting MEK5/ERK5^[24, 25]^ and RAS/MARK^[26]^ pathways. Similarly, downregulation of miR-31 is also shown to repress ovarian cancer^[27]^, hepatocellular carcinoma^[28]^, prostate cancer^[29]^ and other tumor functions^[24]^. These observations suggest that miR-31 may be an interesting target for treatment of cancer and other diseases.^[30-32]^ Interestingly, no protein binding partners have been identified for pre-miR-31^[33]^, suggesting that the mechanisms for regulating biogenesis may be encoded at the RNA level. We therefore sought to examine the RNA structural features that may contribute to the post-transcriptional regulation of pre-miR-31.

Here, we describe the three-dimensional structure of pre-miR-31 and characterized how the stability of secondary structure elements throughout the pre-miR-31 structure affect Dicer processing. The structure presented in this work is the first full-length pre-miR structure determined and significantly adds to the limited known structures of pre-miRs.^[34, 35]^ We examined how three distinct regions of the pre-miR-31 structure; the dicing site, the apical loop, and a short base paired element (junction region) connecting the apical loop and the dicing site, influenced Dicer binding and processing.

We found that modulating the structure of pre-miR-31 at the dicing site by minimizing the size of the internal loop promoted Dicer processing, while structures containing larger internal loops served to inhibit Dicer processing. Furthermore, we demonstrate that the pre-miR-31 apical loop size serves as another point of regulation. Pre-miR-31 constructs with extended junction regions, which restricted the apical loop size, displayed both weaker binding to Dicer and significantly reduced processing. Whereas pre-miR-31 constructs with large apical loops had wild type (WT)-like levels of binding yet reduced processing. These results suggest that the loop size must be tightly controlled, as too small or too large of an apical loop can inhibit pre-miR-31 maturation. Finally, we found that the junction region functions exquisitely to maximize both high affinity binding and efficient processing. We note differences in the secondary structure models derived from nuclear magnetic resonance (NMR) spectroscopy and chemical probing in this junction region. Rather than viewing these structures as incompatible, we demonstrate that both structures likely exist in a dynamic equilibrium where the base paired junction transiently samples the open conformation. We show that the WT pre-miR-31 structure is optimized to maximize both high affinity binding and high efficiency processing. Our data are consistent with a model in which RNAs can self-regulate their processing in the absence of trans-acting RNA-binding proteins. Recent studies demonstrate the importance of pre-miR structural plasticity in regulating their enzymatic processing.^[19, 23]^ Our research cements the hypothesis that pre-miR structure regulates its maturation process and further informs on structural features necessary for effective short hairpin (sh)RNA design.

## Results

### The secondary structure of FL-pre-miR-31 contains three mismatches in the helical stem and three base pairs in the apical loop

The lowest free energy secondary structure of the 71-nt long full length (FL) pre-miR-31 predicted by the RNAStructure webserver^[36]^ is a hairpin composed of three mismatches (A•A, G•A and C•A) in the stem region, a 1×2 internal loop, and three base pairs formed in the junction region between the internal and apical loops. However, recent *in cell* selective 2ʹ hydroxyl acylation analyzed by primer extension (SHAPE) chemical probing studies^[37]^ revealed that the apical loops of pre-miRs are less structured than predicted in the miRbase.^[38-43]^To evaluate the secondary structure of FL pre-miR-31, we performed *in vitro* dimethyl sulfate mutational profiling with sequencing (DMS-MaPseq) on pre-miR-31. The chemical probing derived topology of the entire stem region including the three mismatches is in complete agreement with prediction (**Fig. 1a, Fig. S1**). However, our *in vitro* chemical probing data suggests that residues within and near the predicted apical loop (A33, A34, C35, A40, A41, C42, and C43) are highly reactive, consistent with these residues being unpaired and forming a large, open apical loop structure (**Fig. 1a, Fig. S1**). This is strikingly different from the predicted lowest free energy secondary structure.

**Figure 1.**
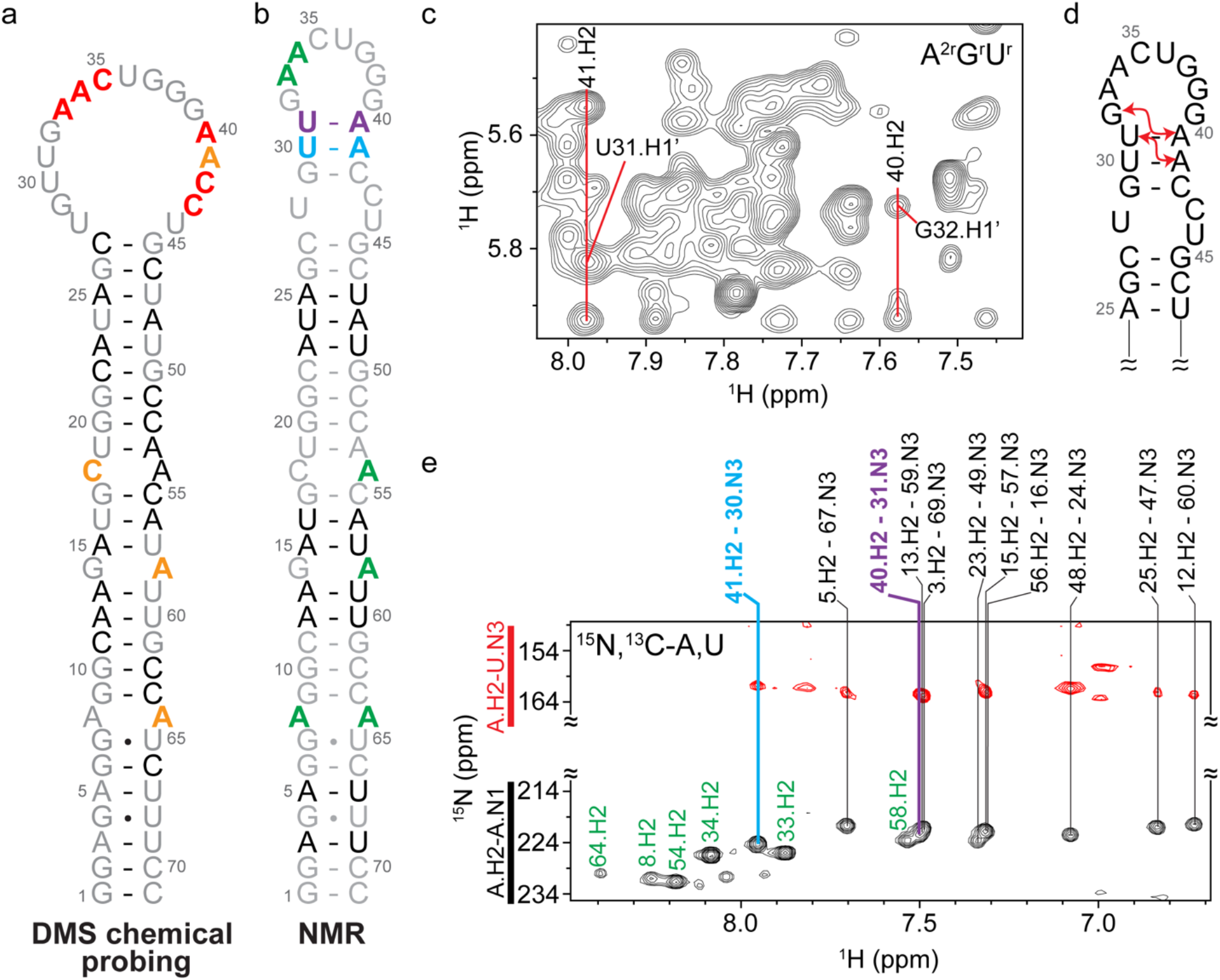
Conflicting secondary structure models for pre-miR-31 apical loop. **a)** Secondary structure derived from *in vitro* DMS-MapSeq where coloring denotes reactivity of given bases. Red=high reactivity, orange=medium reactivity, black=low reactivity, gray=no data available. **b)** Secondary structure derived from NMR characterization. Coloring is based on identification of A-U base pairs (see panel **e**). **c)** Portion of a 2D ^1^H-^1^H NOESY spectrum of an A^2r^G^r^U^r^-labeled FL pre-miR-31. Adenosine cross-strand NOEs consistent with helical stacking in the junction region are indicated. **d)** Secondary structure of the apical loop region highlighting NOEs noted in **c** with red arrows. **e)** Best-selective long-range HNN-COSY spectrum identifying A-U base pairs within FL pre-miR-31. Black peaks are adenosine H2-N1 correlations, red peaks are adenosine H2-uracil N3 correlations. Vertical lines indicate the detection of A-U base pairs. Unpaired adenosines are denoted in green, A-U base pairs in the stem region are denoted in black, junction A-U base pairs are denoted in cyan and purple.

To better understand the molecular details of the pre-miR-31 hairpin, we determined the solution structure of pre-miR-31 using NMR spectroscopy. We used a divide-and-conquer approach to facilitate resonance assignments of full-length (FL) pre-miR-31(**Fig. S2**). We previously reported chemical shift assignments for two fragments, BottomA and BottomB.^[44]^ We completed chemical shift assignments for two additional oligo fragments, TopA (**Fig. S3**) and Top (**Fig. S4**) to guide assignments of the FL pre-miR-31 RNA. However, the large molecular size of FL pre-miR-31 resulted in a severely crowded spectrum, preventing direct assignments based on the oligo controls. To better resolve the complex 2D ^1^H-^1^H NOESY spectrum of FL pre-miR-31, we employed a deuterium-edited approach^[45-47]^ (**Fig. S5**). The combination of methods allowed for complete assignment of non-exchangeable aromatic and anomeric protons (**Fig. S6**).

The topology of the NMR-derived secondary structure of FL pre-miR-31 (**Fig. 1b**) is consistent with the lowest free energy structure. We were particularly interested in the structural features of the apical loop of FL pre-miR-31. Analysis of the ^1^H-^1^H NOESY spectrum of an A^2r^G^r^U^r^-labeled (adenosine C2 and ribose of adenosine, guanosine and uridine residues are protiated, all other sites deuterated) FL pre-miR-31, revealed strong cross-strand NOEs between A41.H2-U31.H1ʹ and A40.H2-G32.H1ʹ (**Fig. 1 c,d**), consistent with a typical A-helical structure in this region. To further explore the base pairing within FL pre-miR-31 we acquired a best selective long-range HNN-COSY^[48]^, which allows for identification of A-U base pairs via detection on the non-exchangeable adenosine C-2 proton rather than detection of the labile imino proton (**Fig. S7**). Here, we see clear evidence for 9 of the 10 expected A-U base pairs within the stem on pre-miR-31 (**Fig. 1e**). The resonance for A53 is broadened beyond detection at pH = 7.5, likely due to the dynamics of the neighboring C18•A54 mismatch. Furthermore, we observe two additional A.H2-U.N3 signals, which correspond to A41-U30 and A40-U31 base pairs (**Fig. 1e**). While A40 and A41 were highly reactive to DMS, and were therefore predicted to be unpaired, we provide direct evidence of base pairing within the apical loop.

Consistent with the NMR-derived secondary structure, pH titration data show that residues A8, A54, A64 (mismatches in the helical stem), and A34 (apical loop) are unpaired due to their high sensitivity to the changes in the pH value of the sample (**Fig. S8**). In contrast, the changes of chemical shifts of A40 and A41 are notably smaller and resemble those measured for base-paired residues from the stem. Additionally, solvent paramagnetic relaxation enhancement (sPRE) data, which reports on the solvent accessibility of FL pre-miR-31, revealed that G29 and A41 do not show large sPRE values (**Fig. S9**) compared to A33, A34, G37 and G38, which are unpaired in the apical loop. Interestingly, for A40 we observe much higher sPRE value indicating high solvent accessibility of the U31-A40 base pair. These observations suggest that U31-A40 may be a nucleation point for opening the loop based on environmental changes. The sequence of pre-miR-31 is highly conserved in mammals, with mutations or deletions present only in the apical loop region (**Fig. S10**). Collectively, our results support the presence of a short base paired element in the junction below the apical loop.

### Tertiary structure of pre-miR-31

To further our structure-based studies, we determined the three-dimensional structure of FL pre-miR-31 (**Fig. 2, Table S1**). The structure is largely an elongated hairpin structure, with three base pair mismatches within the helical stem. Nuclear Overhauser effect (NOE) data are consistent with A-helical stacking of 29-GUU-31 and 40-AAC-42, with strong NOEs between A41.H2-U31.H1’ and A40.H2-G32.H1’ (**Fig. 1c**). The HNN-COSY (**Fig. 1e**) further defines the base pairing within this region, cinching the apical loop structure and limiting the size of the apical loop to 8 nucleotides. The Dicer processing site resides within a 1×2 internal loop containing U28, C43, and U44 (**Fig. 2d**). U28 and U44 are co-planar and adopt a *cis* Watson-Crick/Watson-Crick wobble geometry with C43 positioned above U44. We observed a strong NOE between A54.H2 and U19.H1ʹ, which positions A54 stacked in an A-helical geometry (**Fig. 2e**). No NOEs were observed linking C18 with neighboring residues, therefore C18 was unrestrained in structure calculations and can sample many conformations (**Fig. 2b**). No defined NOEs were observed connecting A13 with G14. However, aromatic-aromatic and aromatic-anomeric NOEs position G14 stacked under A15. G14 and A58 have the potential to form a cis Watson-Crick/Watson-Crick base pair (**Fig. 2f**). The A8•A64 mismatch is well-defined with sequential and cross-strand NOEs (**Fig. 2g**). The structure was refined using global residual dipolar coupling (RDC) restraints. We observed a strong correlation between experimentally determined and back-calculated residual dipolar couplings, further validating the overall structure (**Fig. S11**).

**Figure 2.**
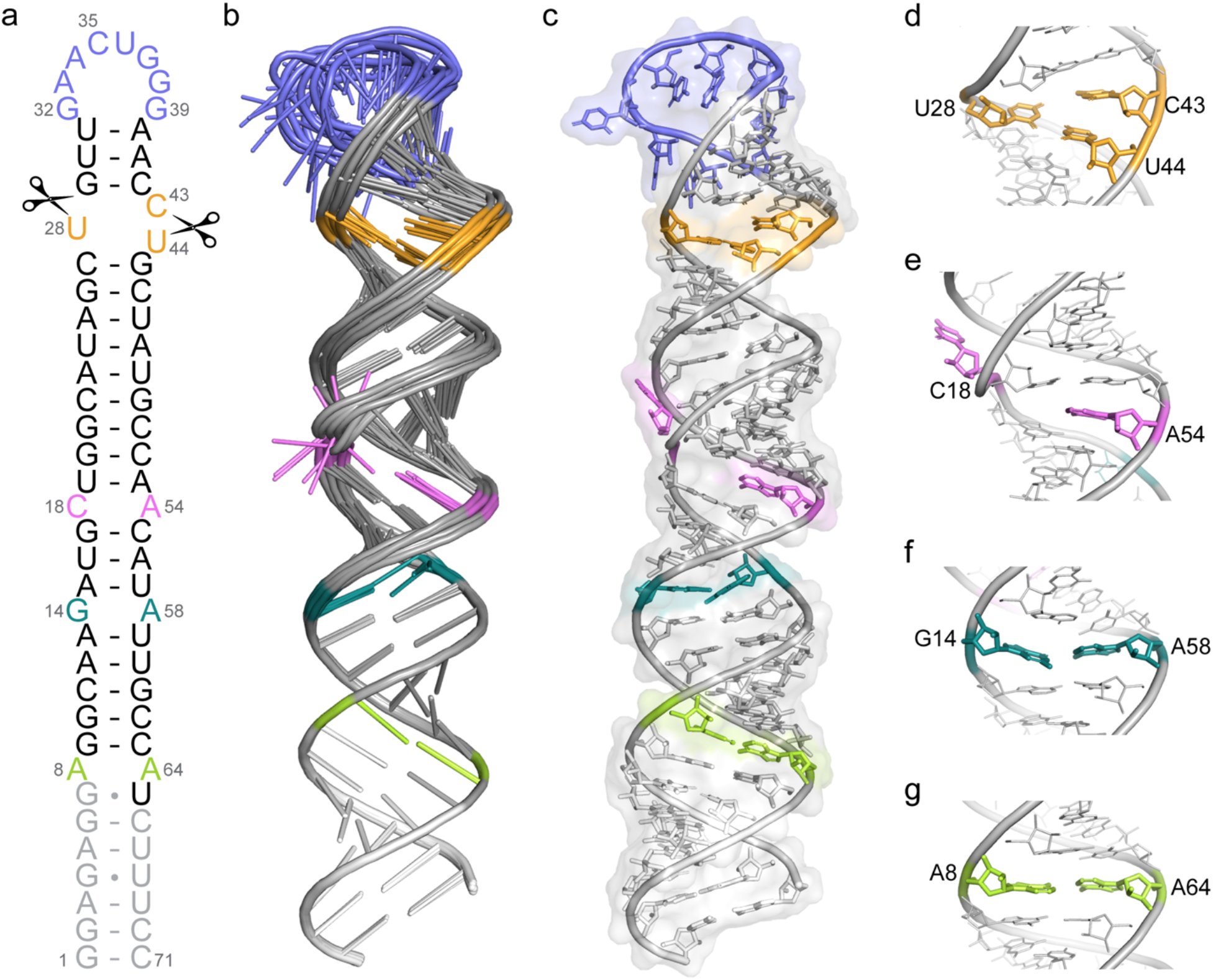
Tertiary structure of pre-miR-31. **a)** NMR-derived secondary structure of FL-pre-miR-31. Dicer cleavage sites are indicated with scissors. Gray nucleotides were included in structural studies but are not present in a Dicing-competent WT pre-miR-31. **b)** Ensemble of 10 lowest energy structures after RDC refinement superimposed over residues 1-13 and 59-71. **c)** Lowest energy structure of pre-miR-31 with a transparent surface rendering. **d)** Enlarged view of the dicing site, colored orange. **e)** Enlarged view of the C•A mismatch, colored pink. **f)** enlarged view of the G•A mismatch, colored teal. **g)** Enlarged view of the A•A mismatch, colored green.

### Mismatches within the helical stem region have no impact on Dicer cleavage

Base pair mismatches are a common feature within the helical stem of precursor microRNAs^[44]^. Increasing the length of the pre-miR helical stem by including additional base paired sequences is detrimental for Dicer processing^[14, 49]^. Studies on fly Dicer-1 suggest that while the length of the pre-miR helical stem is important, the presence of mismatches does not significantly affect Dicer processing^[15]^. However, because pre-miR-31 biogenesis does not appear to be regulated by protein binding partners, we wanted to consider all aspects of pre-miR-31 structure that could be involved in regulating processing. To investigate the role of individual base pair mismatches in the Dicer processing of WT pre-miR-31, we sought to stabilize the G14•A58 mismatch. We made a single point mutation (G14U) which converted the mismatch into a canonical U-A base pair (**Fig. S12**). Quantification of Dicer processing revealed WT-levels of processing of the G14U mutant pre-miR (**Table S2, Fig. S12**).

We previously investigated the pH-dependence of the C18•A54 mismatch and found that A54 is partially protonated at physiological pH, suggesting that these bases can form a C•A^+^ base pair near neutral pH^[44]^. We were therefore interested in testing if mutations that replaced the mismatch with a canonical U-A or C-G base pair (C18U and A54G, respectively) affected the processing by Dicer (**Fig. S12**). As with stabilization of the G•A mismatch, stabilization of the C•A mismatch did not affect the efficiency of Dicer processing (**Table S2, Fig. S12**). We next examined the Dicer processing efficiency of mutant (G14U/A54G) that stabilized both mismatches with canonical base pairs. We found that pre-miR-31 G14U/A54G was processed similarly to WT (**Table S2, Fig. S12**). We next examined the importance of the context of the C•A mismatch by swapping the bases (18ACsw). Again, we observed no significant change in Dicer processing efficiency (**Fig. S12**).

All pre-miR-31 mutant RNAs we examined were cleaved to approximately 90%. Maintaining the same stem length, the absence of one (G14U, C18U, A54G) or two (G14U/A54G) mismatches within the stem of WT pre-miR-31 does not significantly alter the Dicer cleavage efficiency, consistent with studies on fly Dicer-1.^[15]^ However, the measured binding affinity of G14U/A54G for Dicer decreased 2.5-fold relative to WT (**Table S3**). Binding of G14U, C18U, A54G mutants to Dicer were similar to WT while the binding affinity of 18ACsw was slightly enhanced (2-fold). These findings suggest that the mismatches in pre-miR-31 stem are important features for Dicer binding.

### Structure at the cleavage site affects Dicer processing

The RNase III and helicase domains of Dicer interact with the upper stem loop region (which includes the apical loop and the dicing site).^[14, 17, 50]^ Studies strongly indicate that the structure in this region may regulate Dicer processing.^[14, 15, 17, 50]^ To distinguish between the importance of structure at distinct regions within the upper stem loop regions, we employed a mutational approach which reshaped the apical loop and the dicing site, independently.

First, we generated four different Dicer processing site mutants and examined the impact of structure at this site on Dicer processing. We examined two mutations that either minimized (Δ43) or eliminated (Δ43/U44A) the internal loop at the pre-miR-31 Dicer processing site (**Fig. 3a**). The Δ43 construct is processed more efficiently than WT. This is particularly noticeable at timepoints early in the reaction (**Fig. S13**). Interestingly, the Δ43/U44A construct exhibited a slight processing enhancement relative to WT, but was not processed as efficiently as Δ43 (**Fig. 3b**). These findings suggest that a small 1×1 internal loop structure serves as a better substrate for Dicer processing.

**Figure 3.**
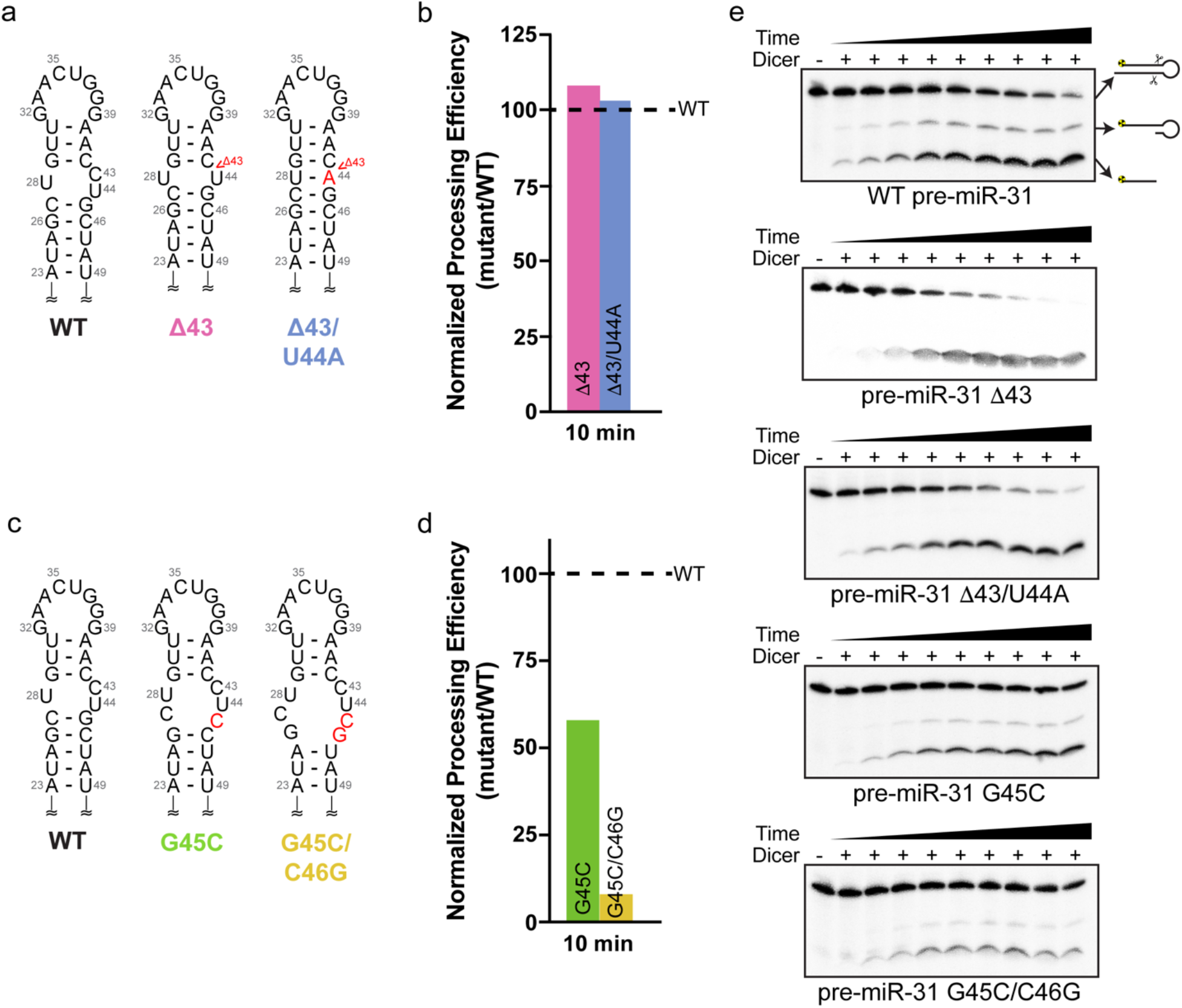
Structure at the dicing site serves as a control element for Dicer processing. **a)** Secondary structures of constructs designed to minimize the internal loop at the dicing site. Mutations are indicated with red lettering. **b)** Dicer processing efficiency for Δ43 and Δ43/U44A mutants normalized to WT pre-miR-31 at 10 min. **c)** Secondary structures of constructs designed to expand the internal loop at the dicing site. Mutations are indicated with red lettering. **d)** Dicer processing efficiency for G45C and G45C/C46G mutants normalized to WT pre-miR-31 at 10 min. **e)** Processing assay gels of hDicer (20 nM) with WT and dicing site mutant pre-miR-31 RNAs (2 nM) at pH = 7.5.

Conversely, we found that mutations that enlarged the internal loop at the dicing site resulted in RNAs that were inefficiently processed by Dicer (**Fig. 3c**). The G45C mutant, which increases the WT 1×2 internal loop to a 2×3 internal loop, has ∼50% reduced processing efficiency while the G45C/C46G mutant (3×4 internal loop) exhibits almost no processing (**Fig. 3d**). Furthermore, we found that Δ43C and Δ43C/U44A, which minimized and eliminated the internal loop, respectively, promoted 5ʹ strand cleavage by Dicer, eliminating the partially processed intermediate, and generating more mature miR (**Fig. 3e**).

Collectively, we found that a 1×1 internal loop at the Dicing site is the best substrate for Dicer processing, while a fully base paired or the native 1×2 internal loop at cleavage site are suboptimal substrates. Pre-miRs with too large of an internal loop around the cleavage site are poor substrates for Dicer to cleave. The binding affinity for Dicer was measured and we found that the Δ43 mutant bound Dicer with near WT affinity, while the Δ43/U44A mutant and the G45C mutant both had a slightly weaker affinity. Introduction of a large internal loop (45/46) reduced binding by ∼6-fold (**Table S3**). Together, our results suggest that Dicer binding affinity and processing efficiency are not strictly correlated, consistent with previous studies^[49]^.

### Size and relative position of the apical loop regulates Dicer processing efficiency and specificity

We next examined the impact or apical loop size on Dicer processing. Apical loop flexibility serves as a control mechanism in many pre-miR/pri-miR elements^[51, 52]^ and the apical loop has been identified as a target for regulation by small molecules or peptides^[53-55]^. Fly Dicer-1 binds to pre-let-7 with 4-nt loop six times weaker than pre-let-7 with 14-nt loop,^[15]^ and the weaker binding leads to poorer cleavage efficiency. However, another study shows that human Dicer binding similarly with different loop sized pre-miR mutants and has uncoupled Dicing activity^[49]^. To further elucidate these findings, we designed two constructs, G32C and G32C/A33C, which minimize the apical loop size by forming one or two canonical base pairs within the otherwise unpaired region (**Fig. 4a**). Dicer binds the G32C RNA (6-nt loop) and the G32C/A33C (4-nt loop) about four times and six times weaker than WT pre-miR-31 (8-nt loop), respectively (**Fig. 4b, Table S3**). The reduced binding affinity correlates with reduced cleavage efficiency (**Fig. 4c**). This result is consistent with observations made with fly Dicer-1^[15]^.

**Figure 4.**
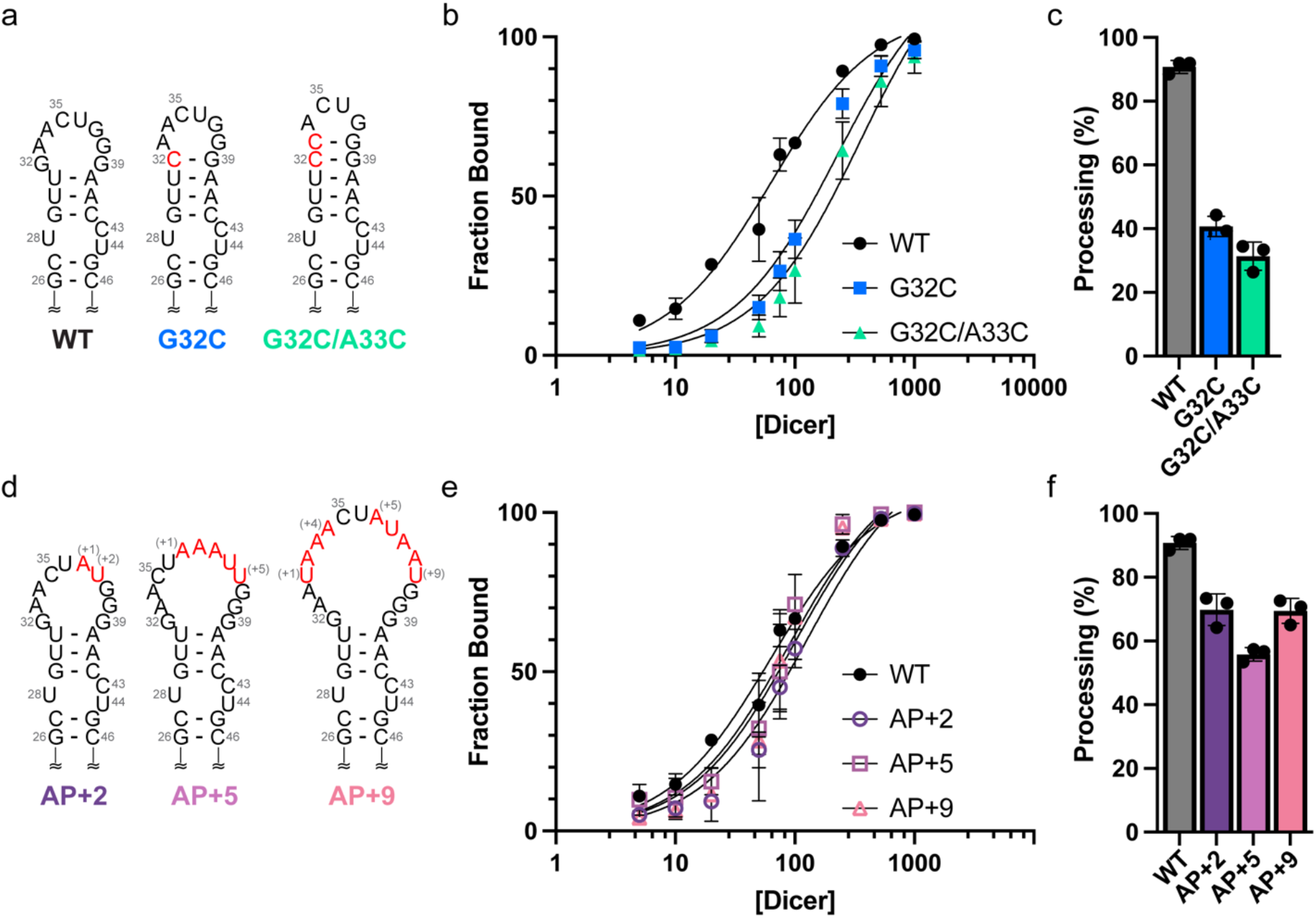
Apical loop size is optimized for efficient Dicer binding and processing. **a)** Secondary structures of constructs designed to minimize the pre-miR-31 apical loop. Mutations are indicated with red lettering. **b)** Quantification of the binding affinity of pre-miR-31 RNAs with Dicer. Solid lines represent best fits to a one site specific binding equation. **c)** Histogram quantifying the Dicer processing efficiencies of pre-miR-31 RNAs at 10 min. **d)** Secondary structures of constructs designed to extend the pre-miR-31 apical loop. Insertions are indicated with red lettering. **e)** Quantification of the binding affinity of pre-miR-31 RNAs with Dicer. Solid lines represent best fits to a one site specific binding equation. **f)** Histogram quantifying the Dicer processing efficiencies of pre-miR-31 RNAs at 10 min. For all binding and processing assays, average and standard deviation from n=3 independent assays are presented. Individual replicates shown with black circles.

Pre-miRs with small apical loops (3-9 nt long) were identified as poor substrates for human Dicer processing, and RNAs with lager apical loops were preferred by Dicer and Drosha^[49]^. We next examined how increasing the apical loop size impacted Dicer cleavage. We added non-native nucleotides to the apical loop regions of pre-miR-31 to generate AP+2 (10-nt loop), AP+5 (13-nt loop) and AP+9 constructs (17-nt loop) (**Fig. 4d**). These larger loop mutants bound human Dicer ∼2-fold weaker than WT (**Fig. 4e, Table S3**). We found that increasing the apical loop size reduced Dicer processing, but not to the same extent as minimizing the apical loop size (**Fig. 4f**).

The reduction in processing efficiency caused by the presence of a larger apical loop can be offset by other factors. Previous studies showed that the apical loop or an internal loop 2-nt from cleavage sites could enhance cleavage efficiency of shRNAs^[17, 21]^. Consistent with previous studies, we observed WT-level processing for a pre-miR-31 construct which contains an 11-nt loop positioned 2-nt from the cleavage site (40UUG, **Fig. S14**). Furthermore, the 40UUG construct generates a U•U mismatch at the dicing site. We demonstrated that dicing site mutants that have 1×1 internal loops at the dicing site are better substrates for Dicer. The restructuring of the Dicing site may further compensate the presence of a larger apical loop.

In addition to enhanced cleavage efficiency, cleavage accuracy is also affected by the loop position. Extension of the helical region between the dicing site and the apical loop results in the generation of mature products of varying lengths. In the G32C/A33C mutant, which shifts the loop position 2-nt up relative to WT, we detected two mature product bands, while for WT, only 1 mature product was observed (**Fig. S15**). We conclude that for pre-miRs, loop size can control Dicer processing efficiency in a bidirectional way. Furthermore, we show that the position of the loop relative to the dicer processing site is essential for accurate and efficient cleavage of Dicer, consistent with the previously described loop counting rule^[17]^.

### Junction residues function as critical control elements for Dicer processing

Our NMR-derived structure of FL pre-miR-31 revealed the presence of three base pairs in a junction region between the apical loop and the dicer cleavage site (**Fig. 1b**). However, *in cell* chemical probing studies revealed that junction residues were highly reactive, suggesting that these base pairs are absent in the presence of Dicer^[37]^. The high reactivity of these nucleotides *in cell* is consistent with our *in vitro* chemical probing studies (**Fig. 1a**) which suggest that pre-miR-31 has a large apical loop region. To resolve these conflicting models, we designed constructs which stabilized or destabilized the junction residues and examined their Dicer binding affinity and Dicer cleavage efficiency.

To mimic the large open loop structure detected by chemical probing, we mutated residues G29, U30, and U31 to prevent base pairing in the junction region (29CAA) (**Fig. 5a**). The processing data for 29CAA reveals that it is a poor substrate for Dicer processing, with only 15% of the 29CAA precursor converted to mature product (**Fig. 5b**). We also designed a construct to stabilize the junction region, as define by NMR data. Here, the junction A-U base pairs were replaced with G-C base pairs (GCclamp, **Fig. 5a**). Interestingly, the GCclamp construct reduced the cleavage efficiency to ∼ 60% (**Fig. 5b**). These data suggest that the stability of the base pairs within the junction region is an important determinant of Dicer processing.

**Figure 5.**
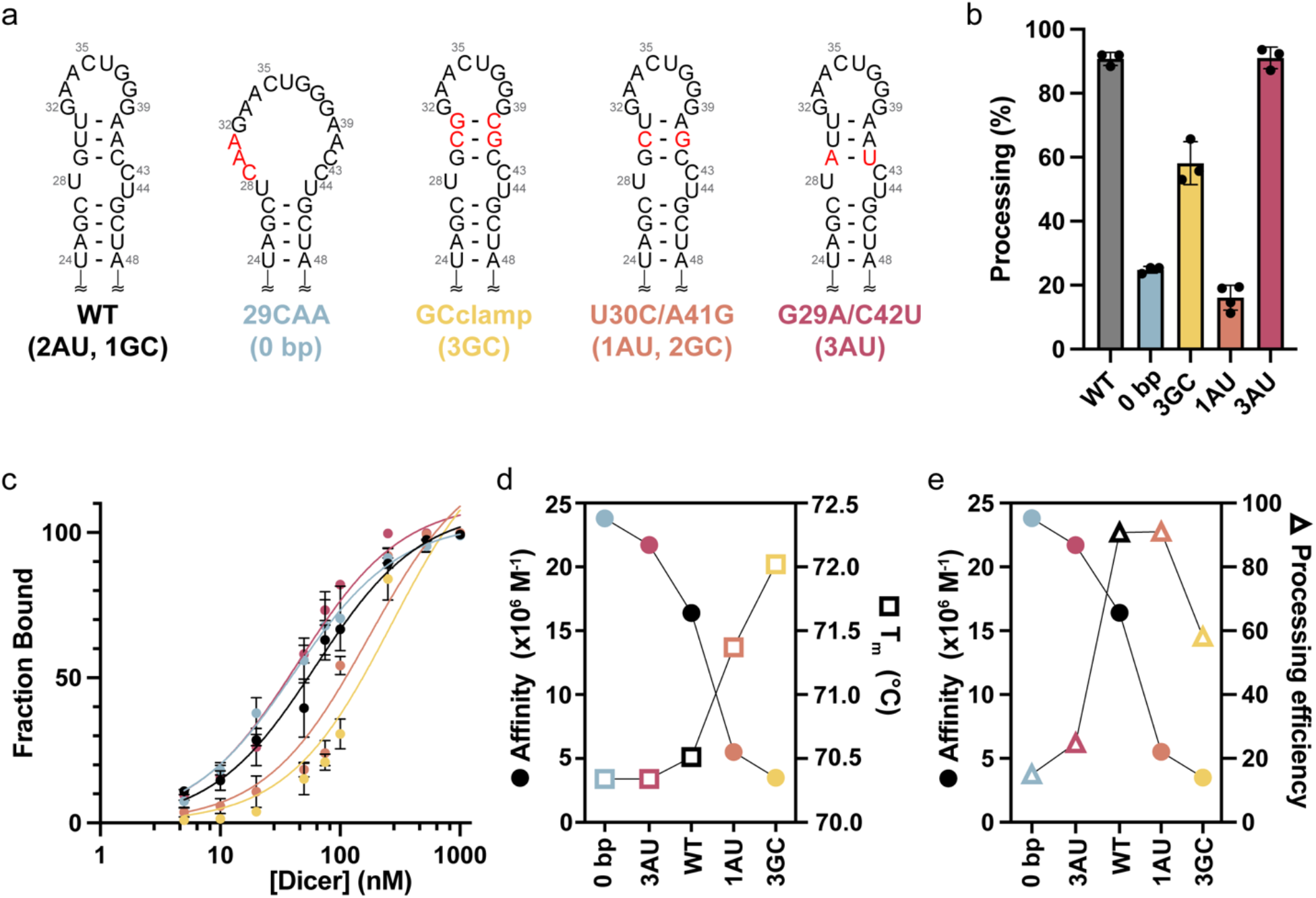
The junction region is a regulatory element within pre-miR-31. **a)** Secondary structures of constructs designed to perturb the stability of the pre-miR-31 junction region. Mutations are indicated with red lettering. **b)** Histogram quantifying the Dicer processing efficiencies of pre-miR-31 RNAs at 10 min. **c)** Quantification of the binding affinity of pre-miR-31 RNAs with Dicer. Solid lines represent best fits to a one site specific binding equation. **d)** Inverse correlation between calculated binding affinity and measured thermal stability (melting temperature, T_m_) for WT and junction region mutations. **e)** Correlation between Dicer binding affinity and Dicer processing efficiency for junction region mutations. For all binding and processing assays, average and standard deviation from n=3 independent assays are presented. Individual replicates shown with black circles.

To further elucidate how the junction stability of pre-miR-31 regulates Dicer processing, we designed two additional junction mutants with different base pairing compositions. The U30C/A41G construct (1AU base pair and 2 GC base pairs, **Fig. 5a**) is processed as efficiently as WT (2AU base pairs, 1 GC base pair) at 10 minutes (**Fig 5b**). Whereas the G29A/C42U mutant (3 AU base pairs, **Fig. 5a**) is processed with ∼20% efficiency (**Fig. 5b**). These data suggest that the stability of the junction region is finely tuned to maximize dicer processing efficiency.

To better characterize the junction stability, we performed thermal denaturation experiments for these constructs. We found that 29CAA, G29A/C42U and WT pre-miR-31 had similar melting temperatures (**Fig. S16, Table S4**), consistent with a model in which they adopt a similar open loop structure. The observed melting temperature of U30C/A41G and GCclamp increased by 1 ℃ and 1.5 ℃, respectively relative to WT pre-miR-31 (**Fig S16, Table S4**). The observed increase in melting temperature suggests that the base pairs in the junction region of these RNAs are more stable than WT.

We show that 29CAA is poorly processed (**Fig. 5b**), however, this RNA adopts an open loop structure, consistent with the Dicer-bound structure identified *in cell*^[37]^. Therefore, we hypothesized that the open loop structure may contribute favorably to dicer binding. We found that 29CAA and G29A/C42U, which both have destabilized junction regions have similar binding affinities, which are slightly tighter than WT (**Fig. 5c, Table S3**). However, mutations that stabilized the junction region (GCclamp, U30C/A41G) exhibited weaker binding relative to wildtype (**Fig. 5C, Table S3**). Collectively, we observe an inverse relationship between junction stability (T_m_) and binding affinity (**Fig. 5d**), consistent with a model in which the binding affinity between Dicer and the pre-miR substrate is determined by the structural stability at the junction.

This delicate balance of structural stability within the junction must be optimized to maximize both high affinity binding and efficient processing. WT pre-miR-31 is precisely tuned to maximize both binding affinity and processing efficiency (**Fig. 5e**). While U30C/A41G maintains high efficiency processing, the increased stabilization of the junction leads to an RNA with reduced binding affinity. Similarly, 29CAA, which has an open loop structure that promotes high affinity binding is poorly processed (**Fig. 5e**).

## Discussion

miRs play an important role in the post-transcriptional regulation of gene expression in cells. miRs are themselves subject to post-transcriptional regulation to ensure appropriate levels of the mature products are produced. Many proteins are known to post-transcriptionally regulate miR biogenesis at either the Drosha and/or Dicer processing steps ^[4, 13, 20]^. While protein-mediated regulation of miR biogenesis can be an important mechanism of control, the intrinsic structural features of pri/pre-miRs can also regulate the enzymatic processing of miRs^[4, 19, 23]^. In fact, in a recent proteomics screen, pre-miR-31 is one of two human miRs (72 miRs examined) with no identified protein binding partners^[33]^. Therefore, we were interested in uncovering the RNA-mediated mechanisms regulating miR-31 biogenesis. To better understand the structural basis for processing, we solved the high-resolution tertiary structure of pre-miR-31. Our structural and biochemical studies provide a framework for optimized design of shRNAs and elucidate distinct mechanisms by which RNA structure helps to regulate Dicer-mediated processing of pre-miR-31 (**Fig. 6**). We found that the presence of mismatches within the pre-miR-31 stem, while a nearly ubiquitous feature of pre-miRs, did not significantly influence the processing of pre-miR-31. We also showed that destabilizing the dicing site by introduction of a larger internal loop inhibited processing of pre-miR-31. Furthermore, we show that apical loop size controls Dicer processing in a bidirectional manner. Finally, we provide strong evidence that stability of pre-miR-31 junction region serves as a potent regulatory factor for Dicer binding and processing.

**Figure 6.**
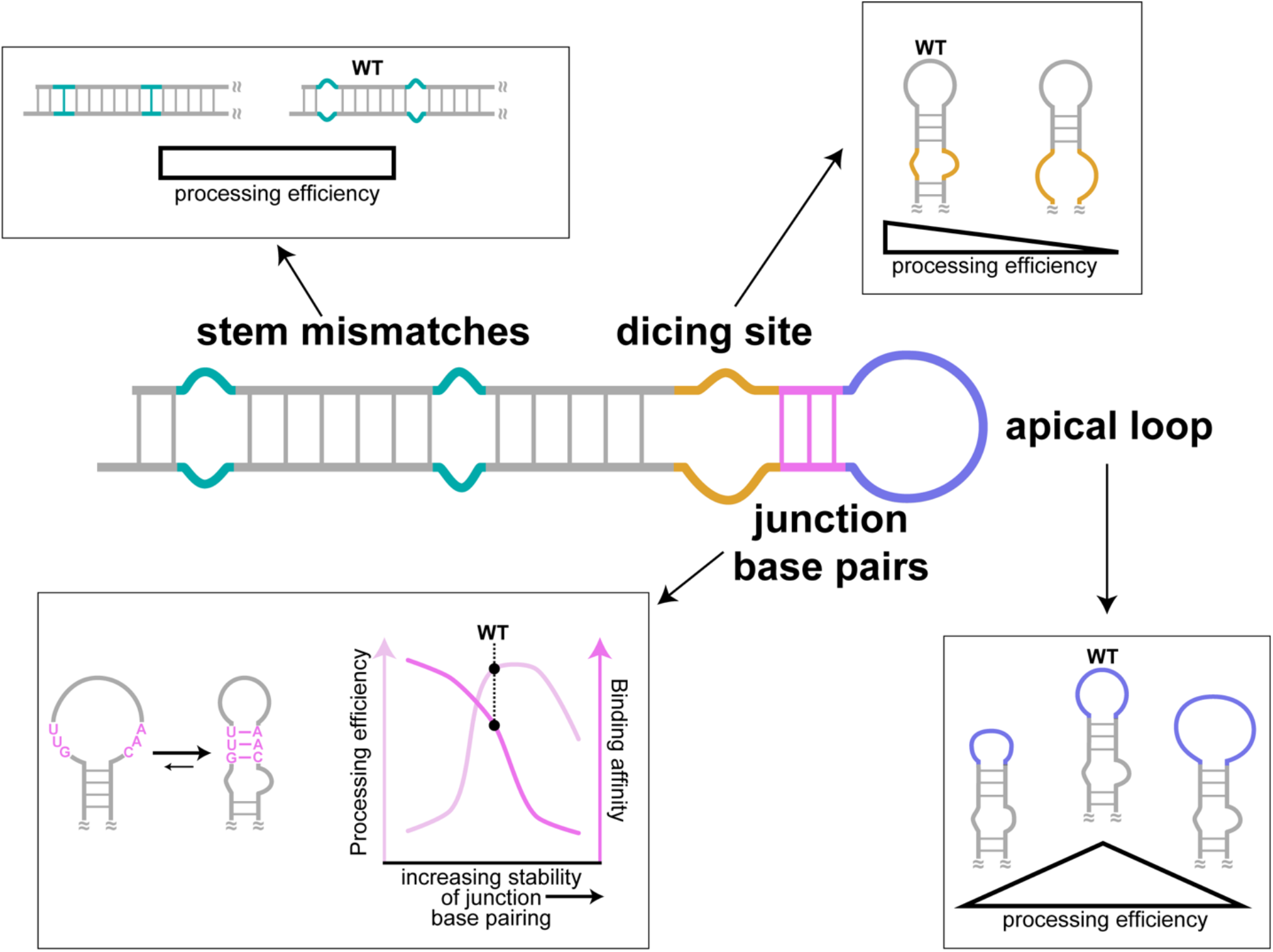
Secondary structure elements and their contribution to the regulation of pre-miR-31 processing. The presence or absence of mismatches within the stem of pre-miR-31 had no impact on Dicer processing. More highly stabilized Dicing sites were processed as efficiently as the WT sequence, but pre-miRs with larger internal loops were not processed efficiently. Similarly, pre-miRs with either too small or too large apical loops were processed less efficiently than WT pre-miR-31. Interestingly, the WT pre-miR-31 has an inherently encoded structural switch at the junction region. Pre-miR-31 appears to sample both an open loop structure, which favors binding, and a closed loop structure, which promotes processing. This allows WT pre-miR-31 to maximize both binding with and processing by Dicer.

Our structure reveals that pre-miR-31 adopts an elongated A-helical structure with three mismatches within the stem region. Both the A•A and G•A mismatches are stacked with their flanking nucleobases. The C•A mismatch is less well-defined. A54 appears to participate in A-helical stacking while C18 samples many conformations. The dicing site is marked by a highly ordered 1×2 internal loop and is linked to the 8-nt apical loop by a 3 base pair junction region.

We previously showed that A54 has an elevated pK_a_ and that a C•A^+^ mismatch within pre-miR-31 can from at near neutral pH^[44]^. A similar pH-regulated conformational switch near the Dicer cleavage site in pre-miR-21 was shown to regulate Dicer processing^[19]^. However, in pre-miR-31, we found that formation of a base pair at the mismatch does not regulate to Dicer processing. In fact, our processing assays show that mutations designed to either stabilize or destabilize the stem mismatches have no effect on Dicer processing. Although the pH-sensitive mismatch within the stem of pre-miR-31 had no effect on Dicer recognition and processing, this and other mismatches may help to regulate Drosha processing^[56]^.

Previous studies show the importance of secondary structure at the dicing site for Dicer cleavage of shRNA and some pre-miRs.^[21]^ Here, we show that substitution to form a 1×1 internal loop at the Dicing site makes itself a slightly better pre-miR substrate for Dicer processing than the native 1×2 internal loop or fully base paired structure at dicing site. However, increasing the internal loop size negatively impacted Dicer processing. Interestingly, we also found that minimizing or eliminating the internal loop at the dicing site promotes 5ʹ strand cleavage by Dicer and effectively eliminates the partially processed intermediate, converting all processed pre-miR to the mature product.

Both apical loop size and position contribute to the regulation of Dicer and Drosha processing^[14, 15, 17, 49]^. Our findings re-emphasized the efficiency control by loop size and efficiency/accuracy control by loop position and provide new insights. Previous studies demonstrate that the presence of a small apical loop inhibits Dicer cleavage^[15, 49]^. We showed not only that a small apical loop inhibits Dicer processing, but also that large apical loops negatively regulate Dicer processing efficiency. We attribute at least a portion of the reduced processing to the weaker binding to Dicer of pre-miRs with small apical loops. We show that as the distance between the cleavage site and the apical loop increases, the processing accuracy decreases. Furthermore, we found that inclusion of a two base pair spacer between the dicing site and the apical loop compensates for the cleavage inhibition caused by a larger apical loop. These findings further validated the loop counting rule^[17]^ in which Dicer has a higher processing efficiency and accuracy when the dicing site is positioned two base pairs below the apical or an internal loop. Our study reveals that loop size is one property that should be optimized when designing shRNAs where large apical loops can reduce Dicer cleavage.

Importantly, we found that the stability of junction region of pre-miR-31 is an inherent regulatory mechanism. Our NMR-derived secondary structure stands in contrast to one revealed by both *in cell* chemical probing^[37]^ and our own *in vitro* chemical probing studies. Secondary structures reported based on chemical probing adopt a large apical loop region, where the junction residues are not engaged in base pairing. We believe that the differences in the NMR and chemical probing derived structures reflect the likely dynamic nature of the base pairs in the junction region, information which can be obstructed in the chemical probing studies. Early chemical probing studies^[57, 58]^ suggest that in the cell, RNAs are generally less folded than *in vitro*. Consistent with this hypothesis, recent *in cell* selective 2ʹ hydroxyl acylation analyzed by primer extension (SHAPE) chemical probing studies revealed that the apical loops of pre-miRs are less structured than predicted in the miRbase.^[37-43]^ Our structural data are consistent with a model in which base pairs in the junction region are very accessible to the solvent and thus more prone to open, so we believe that both an open and cinched junction region exist in a dynamic equilibrium.

We imagine that these two different pre-miR-31 structures both exist and promote distinct favorable interactions with Dicer. We therefore sought to determine the different contributions from the open loop and cinched junction structures. We first examined mutations designed to stabilize the junction region, favoring a cinched junction, consistent with the NMR-derived structure. We found that mutations which stabilized the junction region reduced Dicer binding affinity yet maintained Dicer cleavage. Conversely, we show that mutations which destabilized the junction region, promoting an open apical loop structure, promote binding to Dicer yet inhibit processing. The open apical loop structure sequesters the Dicer cleavage sites in the loop, which may account for the reduced processing efficiency. Collectively, we found that the stability of the pre-miR-31 junction region is optimized to sample both open and cinched conformations to promote both high affinity binding and high efficiency processing. These findings enrich the understanding of how distinct conformations of pre-miR-31 contribute to Dicer binding and processing.

Our newly resolved 3D structure of pre-miR-31 in its processing-competent conformation and elucidation of its intrinsic regulatory mechanism informs on the important role that pre-miR apical loop plasticity plays in controlling Dicer processing. Our structural and biochemical studies are consistent with proposed models of pre-miR processing based on cryo-EM structures of human Dicer^[59]^ and fly Dicer-1^[60]^ bound with pre-miRs. The pre-let-7 bound human Dicer structure revealed that the pre-let-7 RNA adopts multiple conformations^[59]^. In the “pre-dicing state,” Wang and co-workers posit that the pre-let-7 RNA first binds before the structure is adjusted to form a more stable stem^[59]^. This hypothesis is consistent with our findings that the pre-miR-31 large apical loop structure is the preferred substrate for Dicer binding, but that the structure with a cinched junction region is a “dicing-competent” structure. The recent cryo-EM structures of fly Dicer-1 reveal further details of the Dicer-1-pre-miR structure in the “Dicing” state^[60]^. In the “Dicing” structure, the dicing activity of Dicer-1 is inhibited by replacing Mg^2+^ with Ca^2+^. The structure reveals that the pre-miR is highly structured in the “Dicing” state, with the Dicing site sequestered in an A-form helical structure and several base pairs present above the Dicing site. This “Dicing” structure is consistent with our NMR-derived structure, where the stabilization of additional base pairs in the apical loop promotes formation of an extended A-helical structure above the dicing site. Our data suggest that pre-miR-31 is “pre-structured” for Dicer processing. Further structural studies will be necessary to fully-characterize the structural changes in both the pre-miR and Dicer throughout the catalytic cycle.

## Methods

### Preparation of recombinant human Dicer

Human Dicer protein was purified as previously described^[61, 62]^ with modifications. Sf9 cells with infected His-tagged Dicer baculovirus is purchased from University of Michigan protein core. The cell pellet was lysed in ice-cold lysis buffer (50 mM Na_2_HPO_4_ pH = 8.0, 300 mM NaCl, 0.5% Triton X-100, 5% glycerol, 0.5 mM tris(2-carboxyethyl) phosphine (TCEP) and 10 mM imidazole) by sonication. The lysate was pelleted by centrifugation at 30,000 x g for 30 min and the supernatant was mixed with 5 mL pre-equilibrated Ni-NTA resin (Qiagen) in a 50 mL falcon tube. After gently rocking for 1 h at 4 °C, the resin was pelleted by centrifugation at 183 x g for 10 min. The resin was washed with 45 mL wash buffer (50 mM Na_2_HPO_4_ pH = 8.0, 300 mM NaCl, 5% glycerol, 0.5 mM TCEP and 20 mM imidazole) 5 times and eluted with elution buffer (50 mM Na_2_HPO_4_ pH = 8.0, 300 mM NaCl, 5% glycerol, 0.5 mM TCEP and 300 mM imidazole). The elutions were dialyzed against dialysis buffer (20 mM Tris pH = 7.5, 100 mM NaCl, 1 mM MgCl_2_, 0.1% Triton X-100, 50% glycerol). Purified protein was stored at -80 ℃ and total protein concentration was determined by Bradford assay (Thermo Fisher Scientific) and the concentration of Dicer was quantified using ImageJ.

### Preparation of DNA templates

DNA templates for oligo RNAs were purchased from Integrated DNA Technologies (**Table S5**). The DNA templates for *in vitro* transcription were created by annealing the DNA oligonucleotides with an oligonucleotide corresponding to the T7 promoter sequence (5ʹ-TAATACGACTCACTATA-3ʹ). Templates were prepared by mixing the desired DNA oligonucleotide (40 µL, 200 µM) with the complementary oligonucleotide to T7 promoter sequence (20 µL, 600 µM) together, boiling for 3 min, and then slowly cooling to room temperature. The annealed template was diluted with water prior to use to produce the partially double-stranded DNA templates at a final concentration approximately 8 µM.

### Preparation of plasmid templates for *in vitro* transcription

The templates for preparation of the extended pre-miR-31 for DMS-MaPseq and FL pre-miR-31 for NMR studies were generated by overlap-extension (OE) polymerase chain reaction (PCR) using EconoTaq PLUS 2x Master Mix (Lucigen) with primers listed in **Tables S6 and S7**. The OE PCR template was digested with EcoRI and BamHI restriction enzymes and inserted into the pUC-19 plasmid. DNA templates for use in *in vitro* transcription reactions were amplified with EconoTaq PLUS 2x Master Mix (Lucigen) using primers UNIV-pUC19_E105 and miR_tail_3buffer_REV (DMS) or miR31_4R (NMR, **Table S8**).

To ensure the native pre-miR-31 used for processing contained homogeneous 5ʹ-AG sequence, of we included a hammerhead (HH) ribozyme 5ʹ of the pre-miR-31 sequence^[63]^. The native pre-miR-31 template, used to make RNA for processing studies, was generated by OE PCR using EconoTaq PLUS 2x Master Mix (Lucigen) with primers listed in **Table S9**. The OE PCR template was digested with EcoRI and BamHI restriction enzymes and inserted into pUC-19 plasmid. The HH-pre-miR-31-HDV plasmid, which was designed to ensure a homogeneous 3ʹ end of the transcript, was generated by inserting HDV ribozyme sequence to 3ʹ end of HH-pre-miR-31 plasmid construct using the Q5 site-directed mutagenesis kit (New England biolabs) with primers HH-miR-31-HDV-mut-F and HH-miR-31-HDV-mut-R (**Table S10**). All subsequent mutations, deletions, and/or insertions were achieved via site-directed mutagenesis (New England Biolabs Q5 site-directed mutagenesis kit) of the HH-pre-miR-31-HDV plasmid with primers listed in **Table S10**. Templates prepared from plasmids were amplified with EconoTaq PLUS 2x Master Mix (Lucigen) using primers UNIV-pUC19_E105 and HDV-AMP-R (**Table S8**). All primers were purchased from Integrated DNA Technologies. Plasmid identity was verified by Sanger sequencing (Eurofins Genomics) using the universal M13REV sequencing primer.

### Preparation of RNA

RNAs were prepared by *in vitro* transcription in 1× transcription buffer [40 mM Tris base, 5 mM dithiothreitol (DTT), 1 mM spermidine and 0.01% Triton-X (pH = 8.5)] with addition of 3–6 mM ribonucleoside triphosphates (NTPs), 10–20 mM magnesium chloride (MgCl2), 30–40 ng/μL DNA template, 0.2 unit/mL yeast inorganic pyrophosphatase (New England Biolabs)^[64]^, ∼15 μM T7 RNA polymerase and 10–20% (*v*/v) dimethyl sulfoxide (DMSO). Reaction mixtures were incubated at 37 °C for 3–4 h, with shaking at 70 rpm, and then quenched using a solution of 7 M urea and 500 mM ethylenediaminetetraacetic acid (EDTA), pH = 8.5. Reactions were boiled for 3 min and then snap cooled in ice water for 3 min. The transcription mixture was loaded onto preparative-scale 10% denaturing polyacrylamide gels for purification. Target RNAs were visualized by UV shadowing and gel slices with RNA were excised. Gel slices were placed into an elutrap electroelution device (The Gel Company) in 1X TBE buffer. RNA was eluted from the gel at constant voltage (120 V) for ∼24 h. The eluted RNA was spin concentrated, washed with 2 M high-purity sodium chloride, and exchanged into water using Amicon-15 Centrifugal Filter Units (Millipore, Sigma). RNA purity was confirmed on 10% analytical denaturing gels. RNA concentration was quantified via UV-Vis absorbance. Sequences for all RNAs is provided in **Table S11**.

### Dimethyl sulfate (DMS) modification of pre-miR-31 RNA

3 μg of pre-miR-31-tail RNA was denatured at 95 ℃ for 1 min and incubated on ice for another 3 min. Refolding buffer (300 mM sodium cacodylate and 6 mM MgCl_2_) was added to reach total volume of 97.5 uL (for the 0% control), 97.5 μL (for 2.5% modified sample) or 95 μL (for 5% modified sample). The RNA was incubated in refolding buffer at 37 ℃ for 40 min. The RNA was treated with either 2.5 µL DMSO (0% DMS), 2.5 μL DMS (2.5% DMS) or 5 μL DMS (5% DMS) followed by incubation at 37 ℃ while shaking at 250 rpm for 10 min. 60 μL β-mercaptoethanol was added to each reaction to neutralize the residual DMS. The modified RNA was purified using RNA Clean and Concentrator-5 kit (Zymo) according to manufacturer’s instructions.

### RT–PCR with DMS-modified RNA

The methylated RNA was reverse transcribed as follows. 0.2 μM DMS-modified RNA, 2 μl 5× first strand buffer (ThermoFisher Scientific), 1 μl 10 μM reverse primer (miR_tail_RT, **Table S6**), 1 μl dNTP, 0.5 μl 0.1 M DTT, 0.5 μl RNaseOUT and 0.5 μl thermostable group II intron reverse transcriptase, 3^rd^ generation (TGIRT-III, Ingex) were mixed. The mixture was incubated at 57 ℃ for 30 min. After the 30 min incubation, the temperature was increased to 85 ℃ for 5 min. 1 μL RNase H (New England Biolabs) was added to the mixture to digest the RNA. The reverse-transcribed DNA was PCR amplified using Phusion (NEB) for 27 cycles according to the manufacturer’s instruction using primers miR31_buffer_F and miR_tail_RT (**Table S6**). The PCR product was purified by GeneJET PCR purification kit (ThermoFisher Scientific).

### DMS-MaPseq of pre-miR-31 RNA

Illumina sequencing adapters were added by ligation mediated PCR using the NEBNext UltraII DNA Library Prep Kit (New England BioLabs). The libraries were Bioanalyzed on a high sensitivity DNA chip, size selected and sequenced on Illumina Miseq 600 cycles (300×300 paired end). The resulting sequencing reads were adapter trimmed using Trim Galore and aligned using bowtie2 (“bowtie2 --local --no-unal --no-discordant --no-mixed --phred33 40 -L 12”). Each read was compared to its reference sequence to count how many mutations occurred at each nucleotide. All sequencing reads were combined together to calculate the average mutations per base and create a mutational profile.

### Isotopic labeling of RNAs for NMR

Isotopically-labeled RNAs were produced as described above by replacing the rNTP mixture with rNTPs of appropriate isotope labeling. ^15^N/^13^C rNTPs were obtained from Cambridge Isotope Laboratories (CIL, Andover, MA). The partially- and per-deuterated rNTPs used for *in vitro* transcription were obtained from Cambridge Isotope Laboratories (CIL, Andover, MA) or generated in house, as described below. Protiation at the C8 position of perdeuterated rGTP and rATP was achieved by incubation with triethylamine (TEA, 5 equiv) in H_2_O (60 °C for 24 h and for 5 days, respectively). Deuteration of the C8 position of fully protiated GTP and ATP was achieved by analogous treatment with D_2_O (99.8% deuteration; CIL). TEA was subsequently removed by lyophilization.

### NMR experiments

Samples for NMR experiments of Top, TopA, pre-miR-31 and FL pre-miR-31 were prepared in 300-350 μL 100% D_2_O (99.8% deuteration; CIL) or 10% D_2_O/90% H_2_O, 50 mM K-phosphate buffer (pH 7.5), 1 mM MgCl_2_ of 300-600 μM RNA in Shigemi NMR sample tubes. NMR spectra were collected on 600 and 800 MHz Bruker AVANCE NEO spectrometers equipped with a 5 mm three channel inverse (TCI) cryogenic probe (University of Michigan BioNMR Core). NMR spectra of Top and TopA were recorded at 30°C and of pre-miR-31 and FL pre-miR-31 at 37 °C. The isotopic labeling scheme of FL pre-miR-31 used in specific NMR experiment is indicated in the figure legends. NMR data were processed with NMRFx^[65]^ and analyzed with NMRViewJ^[66]. 1^H chemical shifts were referenced to water and ^13^C chemical shifts were indirectly referenced from the ^1^H chemical shift^[67]^.

The signals of nonexchangeable protons of Top and TopA were assigned based on analysis of 2D ^1^H-^1^H NOESY (τ_m_ = 400 ms), 2D ^1^H-^1^H TOCSY (τ_m_ = 80 ms), and ^1^H-^13^C HMQC spectra. Additionally, the 2D NOESY spectrum (τ_m_ = 400 ms) was recorded for A^H^C^H^-labeled Top RNA (A and C fully protiated, G and U perdeuterated). Non-exchangeable ^1^H assignments of FL pre-miR-31 were obtained from 2D NOESY data (τ_m_ = 400 ms) recorded on fully protiated FL pre-miR-31 and A^2r^G^r^-, A^2r^G^r^U^r^-, A^H^C^H^- and G^H^U^6r^-labeled FL pre-miR-31 (superscripts denote sites of protiation on a given nucleoside, all other sites deuterated). ^1^H-^1^H TOCSY and ^1^H-^13^C HSQC spectra of ^15^N/^13^C AG-labeled FL pre-miR-31 were analyzed to facilitate the assignment. The NMR samples for pH titration were prepared with 300 μM ^15^N AU-labeled FL pre-miR-31 in 10% D_2_O/90% H_2_O, 1 mM MgCl_2_ and 10 mM K-phosphate buffer with pH values 5.8, 6.2, 6.5, 7.0, 7.5 and 8.0.

A best-selective long-range HNN-COSY^[48]^ was recorded to identify AU base pairing in FL pre-miR-31. The spectrum was recorded on 560 μM ^15^N AU-labeled FL pre-miR-31 in 10% D_2_O/90% H_2_O, 50 mM K-phosphate buffer (pH=7.5) and 1 mM MgCl_2_. 64 complex points were recorded with a sweep width of 7.4 kHz for ^15^N, and 2048 complex points with a sweep width of 16.6 kHz for ^1^H, 1368 scans per complex increment at 37 °C and 800 MHz.

NMR solvent paramagnetic relaxation enhancement (sPRE)^[68]^, data of FL pre-miR-31 were obtained by measuring R1 relaxation rates^[69]^ as a function of the concentration of paramagnetic compound Gd(DTPA-BMA)^[70]^. We acquired ^1^H-^13^C HSQC-based pseudo-3D experiments at 0.0, 0.8, 1.6, 2.4, 3.2 and 4.8 mM concentration of the paramagnetic compound. Data were acquired on sample containing 480 μM ^15^N/^13^C A,G-labeled FL pre-miR-31 in 100% D_2_O (99.8% deuteration; CIL), 50 mM K-phosphate buffer (pD=7.5) and 1 mM MgCl_2_ at 800 MHz using nine delays (0.02-2s) with two repetitions at every titration point. The data were processed and analyzed using NMRFx^[65]^. The sPRE values were obtained from the peak intensities of well-resolved peaks in the ^1^H-^13^C HSQC-based pseudo-3D experiments. These intensities were fitted to an exponential function (equation 1)^[69]^

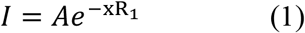

where I is the intensity of the peak, A is the amplitude of the relaxation and R_1_is the longitudinal proton relaxation rate. The sPRE values were obtained from the R_1_ rates determined in the presence of different concentrations of paramagnetic compound Gd(DTPA-BMA) (equation 2)^[68]^

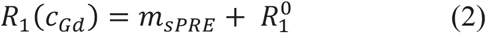

where R_1_(c_Gd_) is the R_1_ measured at the concentration of the paramagnetic compound (c_Gd_), the slope ms_PRE_ corresponds to the sPRE and 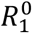 is the fitted R_1_ in the absence of the paramagnetic compound. The error of the sPRE value Δm_sPRE_ were obtained from the linear regression as described previously^[68]^.

We measured ^1^H-^13^C RDCs using IPAP-HSQC experiments^[71]^ for ^15^N/^13^C AG-labeled FL pre-miR-31. Two samples were prepared, an isotropic sample containing 400 μM RNA in 90% H_2_O/10% D_2_O, 50 mM K-phosphate buffer (pH=7.5) and 1 mM MgCl_2_, and an anisotropic sample containing 600 μM FL pre-miR-31 in the same solvent but also including 10 mg/mL Pf1 phage, yielding a solvent ^2^H quadrupole splitting of 11 Hz. 110 complex points were recorded with a sweep width of 8 kHz for ^13^C, and 32768 complex points with a sweep width of 14.7 kHz for ^1^H, 200 scans per complex increment at 800 MHz. Spectra were processed and analyzed with Bruker Topspin.

### Structure calculations

CYANA was used to generate 640 initial structures via simulated annealing molecular dynamics calculations over 128,000 steps. Upper limits for the NOE distance restraints generally set at 5.0 Å for weak, 3.3 Å for medium, and 2.7 Å for strong signals, based on peak intensity. Notable exceptions included intraresidue NOEs between H6/H8 and H2ʹ (4.0 Å) and H3ʹ (3.0 Å). For very weak signals, 6.0 Å upper limit restraints were used, including for sequential H1ʹ-H1ʹ NOEs and intraresidue H5-H1ʹ NOEs. Standard torsion angle restraints were included for regions with A-helical geometry, allowing for ± 25° deviations from ideality (ζ=−73°, α=−62°, β=180°, γ=48°, δ=83°, ɛ=−152°). Torsion angles for mismatches were further relaxed to allow for ± 75° deviation from ideality. Hydrogen bonding restraints were included for experimentally validated base pairs as were standard planarity restraints. Cross-strand P–P distance restraints were employed for A-form helical regions to prevent the generation of structures with collapsed major grooves.^[72]^ A grid search was performed over a broad range of tensor magnitude and rhombicity with weighting of the experimentally determined ^1^H-^13^C residual dipolar couplings (RDCs) constraints. 40 input structures were further minimized after singular value decomposition fits of the RDC weights.

The top 20 CYANA-derived structures were then subjected to molecular dynamics simulations and energy minimization with AMBER.^[73]^ Only upper limit NOE, hydrogen bond, and dipolar coupling restraints were used, along with restraints to enforce planarity of aromatic residues and standard atomic covalent geometries and chiralities.^[72, 74]^ Backbone torsion angle and phosphate-phosphate restraints were excluded during AMBER refinement. Calculations were performed using the RNA.OL3^[75]^ and generalized Born^[76]^ force fields. NMR restraints and structure statistics are presented in **Table S1**.

### ^32^P labeling of RNA

The 5ʹ-end labeling of RNA was performed using 5 pmol of RNA, 1 μL γ-^32^P-ATP (PerkinElmer) and 10 U T4 polynucleotide kinase (New England Biolabs) in a final volume of 10 µL. Before labeling, RNA was boiled for 3 minutes, and snap cooled by placing on ice for another 3 minutes. The radiolabeled RNA was purified on a G-25 column (Cytiva) according to the manufacturer’s instructions. The radiolabeled RNA concentration was determined based on a standard curve which was obtained from the counts per minute of the γ-^32^P-ATP source.

### Dicer processing assay

Human Dicer protein processing assay was performed as previously described with minimal modifications^[19]^. Concentrated recombinant human Dicer protein was diluted in 1X Dicing buffer (24 mM HEPES or 24 mM Bis-Tris, pH 7.5, 100 mM NaCl, 5 mM MgCl_2_, 4 μM EDTA). Dicer enzyme was pre-mixed with 80 U RNaseOUT Recombinant Ribonuclease Inhibitor (Thermo Fisher Scientific) and 5X dicing buffer (120 mM HEPES or 120 mM Bis-Tris, pH 7.5, 0.5 M NaCl, 25 mM MgCl_2_, 0.02 mM EDTA). The ^32^P-labeled RNA was heated to 95℃ for 3 min and then placed on ice for another 3 min. The RNA (1 μL) was added to pre-mixed solution (9 μL) and incubated at 37℃. The final RNA and enzyme concentration are 2 nM and 20 nM, respectively. The reaction is quenched by adding 10 μL quench buffer (98% Formamide, 20 mM EDTA, trace bromophenol blue and xylene cyanol) at 20, 40, 60, 120, 180, 240, 300, 420 and 600 sec respectively. After sample was run on a 12% denaturing polyacrylamide gel, the gel was exposed to a phosphor screen, which was scanned by a Typhoon Phosphor Imager (GE Healthcare). The gel image was quantified analyzed by ImageJ. The Dicer cleavage ratio was calculated as the sum of the intensity of products and partially digested products divided by the sum of the intensity of the products, partially digested products, and remaining substrate. Experiments were performed in triplicate. The average, and standard deviation of the measurements are reported.

### Electrophoretic mobility shift assay

Varied amount of Human recombinant Dicer (5 nM, 10 nM, 20 nM, 50 nM, 75 nM, 100 nM, 250 nM, 525 nM and 1 µM) was incubated with ^32^P-labeled RNA in 24 mM HEPES (pH 7.5), 100 mM NaCl, 5 mM CaCl_2_ on ice for 40 minutes. 5 μL 50% glycerol with trace bromophenol blue and xylene cyanol was added to the mixture and samples were run at 6% native polyacrylamide gel. Then the gel was dried using a gel drying kit (Promega) and exposed to a phosphor screen (overnight). The screen was scanned on a Typhoon Phosphor Imager (GE Healthcare) and quantified and analyzed using ImageJ. Binding ratio was calculated as the intensity of the shifted RNA divided by the intensity of the free RNA and shifted RNA^[49]^. The data were analyzed using equation 3:

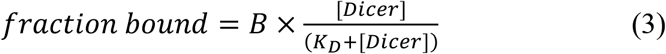

where B is the amplitude of the binding curve^[77]^. Experiments were performed in triplicate. The average, and standard deviation of the measurements are reported.

### CD-thermal denaturation of RNA and data analysis

CD-thermal denaturing of RNAs were performed on JASCO J1500CD spectrometer with a heating rate of 1 ℃ per min from the 5 ℃ to 95 ℃. Data points were collected every 0.5 ℃ with absorbance detection at 260 nm. 20 μM RNA samples were premixed in potassium phosphate buffer (pH=7.5) with 1 mM MgCl_2_. The single transition unfolding melting profiles were analyzed using a two-state model using sloping baselines (equation 4)^[78]^.

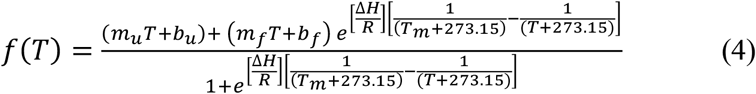

where m_u_ and m_f_ are the slopes of the lower (unfolded) and upper (folded) baselines, b_u_ and b_f_ are the y-intercepts of the lower and upper baselines, respectively. ΔH (in kcal/mol) is the enthalpy of folding and T_m_ (in °C) is the melting temperature, R is the gas constant (0.001987 kcal/(Kmol)). Experiments were performed in triplicate. The average, and standard deviation of the measurements are reported.

## Abbreviations

(miR): microRNA
(pri-miR): primary microRNA
(pre-miR): precursor microRNA
(DGCR8): DiGeorge syndrome critical region 8
(GTP): guanosine triphosphate
(nt): nucleotide
(Ago): Argonaute
(ILF3): Interleukin enhancer-binding factor 3
(RBP): RNA binding protein
(TUTase): terminal uridyltransferase
(CRC): colorectal cancer
(MEK5): extracellular signal-regulated kinase
(ERK5): extracellular-regulated protein kinase 5
(MARK): mitogen-activated protein kinase
(miR-31): microRNA-31
(NMR): nuclear magnetic resonance
(shRNA): short hairpin RNA
(WT): wild type
(FL): full length
(NOESY): nuclear Overhauser effect spectroscopy
(COSY): correlated spectroscopy
(SHAPE): selective 2ʹ hydroxyl acylation analyzed by primer extension
(DMS-MaPseq): dimethyl sulfate mutational profiling with sequencing
(NOE): nuclear Overhauser effect
(sPRE): solvent paramagnetic relaxation enhancement
(RDC): residual dipolar coupling

## Data availability

Resonance assignments have been deposited in the BMRB (miR-31_TopA: 51697, miR-31_Top: 51698, pre-miR-31: 31061). NMR-derived structures have been deposited in the PDB (pre-miR-31: 8FCS). Fastq files were deposited in Gene Expression Omnibus (GEO), accession number pending.

## Conflicts of interest

The authors declare that they have no conflict of interest.

## Acknowledgements

This work was supported by National Institute of General Medical Sciences of the National Institutes of Health grant R35 GM138279 (to S.C.K.), Research Corporation for Science Advancement Cottrell Scholar Award 28248 (S.C.K.), and the Pew Charitable Trusts Scholars Program (S.C.K.). Research reported in this publication was supported by the University of Michigan BioNMR Core Facility (U-M BioNMR). U-M BioNMR Core is grateful for support from U-M including the College of Literature, Sciences and Arts, Life Sciences Institute, College of Pharmacy and the Medical School along with the U-M Biosciences Initiative.

## Notes

### Competing Interest Statement

The authors have declared no competing interest.

