## Supplement for "Structure of pre-miR-31 reveals an active role in Dicer processing"

**Table S1. NMR restraints and structural statistics for the FL pre-miR-31 structure.[1]**

| | |
|---|---|
| **Cyana[2]** | |
| NOE-derived restraints | 516 |
| Intraresidue | 138 |
| Sequential | 359 |
| Long range (\|i – j\| > 1) | 19 |
| H-bond restraints | 134 |
| RDC | 30 |
| NOE restraints/residue | 7.3 |
| Total restraints/residue | 9.6 |
| Target function (Å2 ) | 1.99 ± 0.02 |
| Upper distance viol. (Å2 ) | 0.0269 ± 0.0002 |
| Lower distance viol. (Å2 ) | 0.0412 ± 0.0001 |
| RMSD[3] (Å) | 2.68 ± 0.15 |
| Q value | 5.4 ± 0.3 % |
| **Amber[4]** | |
| Amber energy | -16,427.7 |
| Distance | 160.1 |
| Torsion | 7.3 |
| RMSD[3] (Å) | 2.16 ± 0.48 |
| RMSD lower stem (1-13, 59-71) (Å) | 0.22 ± 0.08 |
| RMSD full stem (1-27, 45-71) (Å) | 1.30 ± 0.37 |
| RMSD bp in loop (29-31, 40-42) (Å) | 0.33 ± 0.11 |
| RMSD loop (32-39) (Å) | 3.23 ± 1.04 |
| **MolProbity analysis[5]** | |
| Clashscore | 0.44 |
| Probably wrong sugar pucker (%) | 0 |
| Bad backbone conformation (%) | 4 |
| Bad bonds (%) | 0 |
| Bad angles (%) | 0 |

[1]Statistics are reported for the entire structure unless otherwise specified.
[2]Statistics for the 20 structures with lowest target function
[3]RMSD: root mean squared deviation.
[4]Statistics for the 20 lowest energy structures
[5]The 20 amber-refined structures were evaluated using the MolProbity webserver[1, 2]

**Table S2. Dicer cleavage of pre-miR-31 mutations at 10 minutes.**

| Region mutated | RNA constructs | Cleavage efficiency (%)[a] |
|---|---|---|
| - | pre-miR-31 WT | 90.8 ± 1.7 |
| Stem mutations | pre-miR-31 G14U | 87.6 ± 2.7 |
| | pre-miR-31 C18U | 83.8 ± 6.1 |
| | pre-miR-31 A54G | 82.2 ± 3.2 |
| | pre-miR-31 G14U/A54G | 88.0 ± 6.4 |
| | pre-miR-31 18ACsw | 83.4 ± 1.4 |
| Dicing site mutations | pre-miR-31 Δ43C | 98.0 ± 2.3 |
| | pre-miR-31 Δ43/U44A | 93.5 ± 1.2 |
| | pre-miR-31 G45C | 52.2 ± 4.8 |
| | pre-miR-31 G45C/C46G | 7.0 ± 4.9 |
| Apical loop mutations | pre-miR-31 G32C | 41.8 ± 3.2 |
| | pre-miR-31 G32C/A33C | 31.4 ± 4.4 |
| | pre-miR-31 AP+2 | 69.8 ± 5.0 |
| | pre-miR-31 AP+5 | 55.8 ± 2.2 |
| | pre-miR-31 AP+9 | 69.5 ± 3.9 |
| | pre-miR-31 40UUG | 91.0 ± 3.2 |
| Junction mutations | pre-miR-31 29CAA | 25 ± 5 |
| | pre-miR-31 3GCclamp | 4 ± 1 |
| | pre-miR-31 U30C A41G | 6 ± 2 |
| | pre-miR-31 G29A C42U | 22 ± 3 |

[a] Average and standard deviation from n=3 independent assays are presented.

**Table S3. Dicer binding affinity of pre-miR-31 mutations.**

| Region mutated | RNA constructs | Binding affinity (x$10^6$ $M^{-1}$)[a] |
|---|---|---|
| - | pre-miR-31 WT | 17 ± 2 |
| Stem mutations | pre-miR-31 G14U | 19 ± 3 |
| | pre-miR-31 C18U | 13 ± 1 |
| | pre-miR-31 A54G | 23 ± 4 |
| | pre-miR-31 G14U/A54G | 6 ± 1 |
| | pre-miR-31 18ACsw | 36 ± 6 |
| Dicing site mutations | pre-miR-31 Δ43 | 11 ± 2 |
| | pre-miR-31 Δ43/U44A | 7 ± 1 |
| | pre-miR-31 G45C | 9 ± 2 |
| | pre-miR-31 G45C/C46G | 3 ± 1 |
| Apical loop mutations | pre-miR-31 G32C | 5 ± 1 |
| | pre-miR-31 G32C/A33C | 3 ± 1 |
| | pre-miR-31 AP+2 | 8 ± 2 |
| | pre-miR-31 AP+5 | 11 ± 3 |
| | pre-miR-31 AP+9 | 11 ± 3 |
| | pre-miR-31 40UUG | 18 ± 2 |
| Junction mutations | pre-miR-31 29CAA | 25 ± 5 |
| | pre-miR-31 3GCclamp | 4 ± 1 |
| | pre-miR-31 U30C A41G | 6 ± 2 |
| | pre-miR-31 G29A C42U | 22 ± 3 |

[a] Average and standard deviation from n=3 independent assays are presented.

**Table S4. Thermal stability of pre-miR-31 RNAs.**

| Region mutated | RNA constructs | $T_m$ (°C)[a] |
|---|---|---|
| - | pre-miR-31 WT | 70.4 ± 0.3 |
| Junction mutations | pre-miR-31 29CAA | 70.3 ± 0.2 |
| | pre-miR-31 3GCclamp | 72.0 ± 0.2 |
| | pre-miR-31 U30C A41G | 71.4 ± 0.3 |
| | pre-miR-31 G29A C42U | 70.3 ± 0.2 |

[a] $T_m$ values were obtained by fitting CD thermal denaturation profiles to a two-state unfolding model using sloping baselines.

**Table S5. Synthetic DNA templates and associated RNA constructs.**

| | 5'-sequence-3'[a,b,c] | |
|---|---|---|
| | **DNA** | **RNA** |
| Top | mGmGCATAGCAGGTTCCCAGTTCAACAGCTATGCC*TATAGTGAGTCGTATTA* | GGCAUAGCUGUUGAACUGGGAACCUGCUAUGCC |
| TopA | mGmGCATAGCCGTAGCTATGCC*TATAGTGAGTCGTATTA* | GGCAUAGCUACGGCUAUGCC |

[a] m denotes 2′-*O*-Me modification of the primer.
[b] Italicized nucleotides correspond to the sequence complementary to the T7 promoter.
[c] Red nucleotides indicate non-native tetraloop sequences.

**Table S6. DNA primers for pre-miR-31-tail (DMS) experiments.**

| OE-PCR primers | 5'-sequence-3'[a] |
|---|---|
| miR31_tail-1F | GCAGCTGAATTCTTCTAATACGACTCACTATAGGAGACCTCGAGTAGAGGTCAAAA |
| miR31_tail-2R | CCAGCATCTTGCCTCCTCTCCTTTTGACCTCTACTCGAGGTCTCCTATAGTG |
| miR31_tail-3F | GGAGGCAAGATGCTGGCATAGCTGTTGAACTGGGAACCTGCTATGCCAACAT |
| miR31_tail-4R | AGTTGTTTGGAAAGATGGCAATATGTTGGCATAGCAGGTTCCC |
| miR31_tail-5F | TTGCCATCTTTCCAAACAACTCGAGTAGAGTTGACAACAAAGAAACAACAACAACAACGG |
| miR31_tail-6R | GCAGGAGGATCCGTTGTTGTTGTTGTTTCTTTGTTGTC |
| miR_tail_RT | GTTGTTGTTGTTGTTTCTTTGTTGTCAACTCTACTCGAGTTGTTT |
| miR31_buffer_F | GGAGACCTCGAGTAGAGGTCAAAAGGAGAGG |

**Table S7. DNA primers for generation of the pre-miR-31 (NMR) template.**

| OE-PCR primers | 5'-sequence-3' |
|---|---|
| miR-31FL-OE-1F | GTGTCAGAATTCTAATACGACTCACTATAGGAGAGGAGGCAAG |
| miR-31FL-OE-2R | CCAGTTCAACAGCTATGCCAGCATCTTGCCTCCTCTCCTATAGTGAGTCGTA |
| miR-31FL-OE-3F | GCTGGCATAGCTGTTGAACTGGGAACCTGCTATGCCAACATATTGCCATCTTT |
| miR-31FL-OE-4R | CATAGCGGATCCGGAAAGATGGCAATATGTTGGCATAGCAGGT |

**Table S8. Amplification primers for template.**

| Amplification primers | 5'-sequence-3'[a] | application |
|---|---|---|
| UNIV-pUC19_E105 | TCTTCGCTATTACGCCAGCTGGCGAAA | Forward primer for amplification of DNA template for pre-miR-31 NMR construct and all processing constructs |
| HDV-AMP-R | mUmAATGTGAGAATTGGCTACGTTGAAACAACGCATTACCG | Reverse primer for amplification of DNA template for all pre-miR-31 processing constructs |
| miR31_4R | mGmGAAAGATGGCAATATGTTGGCATAGCAGGTT | Reverse primer for amplification of DNA template for pre-miR-31 NMR construct |
| miR_tail_3buffer_REV | mGmUTGTTGTTGTTGTTTCTTTGTTGTCAACTCTACTCGAGTTG | Reverse primer for amplification of DNA template for pre-miR-31 DMS construct |

[a] m denotes 2′-*O*-Me modification of the primer.

**Table S9. DNA primers for HH-pre-miR-31 template.**

| OE-PCR primers | 5'-sequence-3' |
|---|---|
| HH_miR31_Nat_1F | CCGGAATTCTAATACGACTCACTATAGGGCTC |
| HH_miR31_Nat_2R | ACGTACCCTGATGGTGTACGAGCCCTATAGTGAGTCGTATTA |
| HH_miR31_Nat_3F | ACACCATCAGGGTACGTTTTTCAGACACCATCAGGGTCTGGCATCTTGCCTCT |
| HH_miR31_Nat_4R | CTGACGGTACCGGGTACCGTTTCGTCCTCACGGACTCATCAGAGGCAAGATGCCAGACC |
| HH_miR31_Nat_5F | ACCCGGTACCGTCAGGCAAGATGCTGGCATAGCTGTTGAACTGGGAACCTGCTATGCCAA |
| HH_miR31_Nat_6R | CCGTCGCGGATCCATGGCAATATGTTGGCATAGCAGGTTCCCAGT |

**Table S10. Mutation DNA primers for processing constructs.**

| Mutagenesis primers | 5'-sequence-3' | Application |
|---|---|---|
| HH-miR-31-HDV-mut-F | TAATGCGTTGTTTCAACGTAGCCAATTC TCACATTAGGATCCTCTAGAGTCGAC | Forward primer to insert HDV-like sequence to 3' end of HH-pre-miR-31 |
| HH-miR-31-HDV-mut-R | CCGACACTACGACGGGGACGTTTCTCAC TCAGTGTCATGGCAATATGTTGGCATAG | Reverse primer to insert HDV-like sequence to 3' end of HH-pre-miR-31 |
| HH-A54G-HDV-mut-R | CCGACACTACGACGGGGACGTTTCTCAC TCAGTGTCATGGCAATATGCTGGCATAG | Reverse primer to insert HDV-like sequence to 3' end of HH-pre-miR-31-A54G |
| miR-31-Nat-G14U-Mut-F | CGTCAGGCAATATGCTGGCATAG | Forward primer for G14U construct mutation |
| miR-31-Nat-G14U-Mut-R | GTACCGGGTACCGTTTCG | Reverse primer for G14U construct mutation |
| miR31-G14U-11nt-F | GGTCTGGCATATTGCCTCTGA | Forward primer for G14U hammerhead complementary sequence mutation |
| miR31-G14U-11nt-R | CTGATGGTGTCTGAAAAACG | Reverse primer for G14U hammerhead complementary sequence mutation |
| miR-31-Nat-C18U-Mut-F | AGGCAAGATGTTGGCATAGCTGTTG | Forward primer for C18U construct mutation |
| miR-31-Nat-C18U-Mut-R | GACGGTACCGGGTACCGT | Reverse primer for C18U construct mutation |
| miR31-C18U-11nt-F | TCAGGGTCTGACATCTTGCCT | Forward primer for C18U hammerhead complementary sequence mutation |
| miR31-C18U-11nt-R | TGGTGTCTGAAAAACGTAC | Reverse primer for C18U hammerhead complementary sequence mutation |
| miR31-18Acsw-mut-F | TGCTATGCCACCATATTGCCATG | Forward primer for 18Acsw construct mutation |
| miR31-18Acsw-mut-R | GGTTCCCAGTTCAACAGC | Reverse primer for 18Acsw construct mutation |
| miR-31-Nat-29CAA-Mut-F | TGGCATAGCTCAAGAACTGGGAACC | Forward primer for 29CAA construct mutation |
| miR-31-Nat-29CAA-Mut-R | GCATCTTGCCTGACGGTA | Reverse primer for 29CAA construct mutation |
| miR31-nat-G32C-F | CATAGCTGTTCAACTGGGAACC | Forward primer for G32C construct mutation |
| miR31-nat-G32C-R | CCAGCATCTTGCCTGACG | Reverse primer for G32C construct mutation |
| miR31-Nat-G32C/A33C-F | CATAGCTGTTCCACTGGGAACCTG | Forward primer for G32C/A33C construct mutation |
| miR31-Nat-G32C/A33C-R | CCAGCATCTTGCCTGACG | Reverse primer for G32C/A33C construct mutation |
| miR31-nat-40UUG-mut-F | TTGAACTGGGTTGCTGCTATGCCAAC | Forward primer for 40UUG construct mutation |
| miR31-nat-40UUG-mut-R | CAGCTATGCCAGCATCTTG | Reverse primer for 40UUG construct mutation |
| miR31-nat-A54G-mut-F | TGCTATGCCAGCATATTGCCAT | Forward primer for A54G construct mutation |
| miR31-nat-A54G-mut-R | GGTTCCCAGTTCAACAGC | Reverse primer for A54G construct mutation |

| | | |
|---|---|---|
| miR31-AP+2-mut-F | ATGGGAACCTGCTATGCCAA | Forward primer for AP+2 construct mutation |
| miR31-AP+2-mut-R | AGTTCAACAGCTATGCCAG | Reverse primer for AP+2 construct mutation |
| miR31-AP+5-mut-F | ATAATGGGAACCTGCTATGCCAA | Forward primer for AP+5 construct mutation |
| miR31-AP+5-mut-R | AGTTCAACAGCTATGCCAG | Reverse primer for AP+5 construct mutation |
| miR31-AP+9-mut-F | TATAATGGGAACCTGCTATGCCAA | Forward primer for AP+9 construct mutation |
| miR31-AP+9-mut-R | GTTTATTCAACAGCTATGCCAGCAT | Reverse primer for AP+9 construct mutation |
| miR31-3Gcclamp-mut-F | TGGGCGCCTGCTATGCCAACATATTG | Forward primer for 3Gcclamp construct mutation |
| miR31-3Gcclamp-mut-R | GTTCCGCAGCTATGCCAGCATCTTG | Reverse primer for 3Gcclamp construct mutation |
| miR31-G45C-mut-F | CTGGGAACCTCCTATGCCAAC | Forward primer for G45C construct mutation |
| miR31-G45C-mut-R | TTCAACAGCTATGCCAGC | Reverse primer for G45C construct mutation |
| miR31-G45C C46G-mut-F | CTGGGAACCTCGTATGCCAACATATTG | Forward primer for G45C/C46G construct mutation |
| miR31-G45C C46G-mut-R | TTCAACAGCTATGCCAGC | Reverse primer for G45C/C46G construct mutation |
| miR31-d43C-mut-F | TGCTATGCCAACATATTGC | Forward primer for Δ43 construct mutation |
| miR31-d43C-mut-R | GTTCCCAGTTCAACAGCTATG | Reverse primer for Δ43 construct mutation |
| miR31-4344A-mut-F | AACTGGGAACAGCTATGCCAAC | Forward primer for Δ43/U44A construct mutation |
| miR31-4344A-mut-R | CAACAGCTATGCCAGCAT | Reverse primer for Δ43/U44A construct mutation |
| G29A C42U-mut-FWD | TGGGAATCTGCTATGCCAACATATTG | Forward primer for G29A/C42U construct mutation |
| G29A C42U-mut-REV | GTTCAATAGCTATGCCAGCATCTTG | Reverse primer for G29A/C42U construct mutation |
| U30C A41G-mut-FWD | TGGGAGCCTGCTATGCCAACATATTG | Forward primer for U30C/A41G construct mutation |
| U30C A41G-mut-REV | GTTCAGCAGCTATGCCAGCATCTTG | Reverse primer for U30C/A41G construct mutation |

**Table S11. RNA sequences used for structural and processing studies.**

| construct name | 5'-RNA sequence-3' | Application |
|---|---|---|
| miR-31_DMS | GGAGACCUCGAGUAGAGGUCAAAGGAGAGGAGGC<br>AAGAUGCUGGCAUAGCUGUUGAACUGGGAACCUGC<br>UAUGCCAACAUAUUGCCAUCUUUCCAAACAACUCGA<br>GUAGAGUUGACAACAAAGAAACAACAACAACAAC | DMS chemical probing |
| FL-pre-miR-31 | GGAGAGGAGGCAAGAUGCUGGCAUAGCUGUUGAAC<br>UGGGAACCUGCUAUGCCAACAUAUUGCCAUCUUUCC | Structure |
| WT pre-miR-31 | AGGCAAGAUGCUGGCAUAGCUGUUGAACUGGGAAC<br>CUGCUAUGCCAACAUAUUGCCAU | Processing |
| WT pre-miR-31 -G14U | AGGCAAUAUGCUGGCAUAGCUGUUGAACUGGGAAC<br>CUGCUAUGCCAACAUAUUGCCAU | Processing |
| WT pre-miR-31-C18U | AGGCAAGAUGUUGGCAUAGCUGUUGAACUGGGAAC<br>CUGCUAUGCCAACAUAUUGCCAU | Processing |
| WT pre-miR-31-18Acsw | AGGCAAGAUGAUGGCAUAGCUGUUGAACUGGGAAC<br>CUGCUAUGCCACCAUAUUGCCAU | Processing |
| WT pre-miR-31-A54G | AGGCAAGAUGCUGGCAUAGCUGUUGAACUGGGAAC<br>CUGCUAUGCCAGCAUAUUGCCAU | Processing |
| WT pre-miR-31-40UUG | AGGCAAGAUGCUGGCAUAGCUGUUGAACUGGGUUG<br>CUGCUAUGCCAACAUAUUGCCAU | Processing |
| WT pre-miR-31-29CAA | AGGCAAGAUGCUGGCAUAGCUCAAGAACUGGGAAC<br>CUGCUAUGCCAACAUAUUGCCAU | Processing |
| WT pre-miR-31-G32C | AGGCAAGAUGCUGGCAUAGCUGUUCAACUGGGAAC<br>CUGCUAUGCCAACAUAUUGCCAU | Processing |
| WT pre-miR-31-G32C/A33C | AGGCAAGAUGCUGGCAUAGCUGUUCCACUGGGAAC<br>CUGCUAUGCCAACAUAUUGCCAU | Processing |
| WT pre-miR-31-G14U/A54G | AGGCAAUAUGCUGGCAUAGCUGUUGAACUGGGAAC<br>CUGCUAUGCCAGCAUAUUGCCAU | Processing |
| WT pre-miR-31-AP+2 | AGGCAAGAUGCUGGCAUAGCUGUUGAACUAUGGGA<br>ACCUGCUAUGCCAACAUAUUGCCAU | Processing |
| WT pre-miR-31-AP+5 | AGGCAAGAUGCUGGCAUAGCUGUUGAACUAUAAUG<br>GGAACCUGCUAUGCCAACAUAUUGCCAU | Processing |
| WT pre-miR-31-AP+9 | AGGCAAGAUGCUGGCAUAGCUGUUGAAUAAACUAU<br>AAUGGGAACCUGCUAUGCCAACAUAUUGCCAU | Processing |
| WT -miR-31-3Gcclamp | AGGCAAGAUGCUGGCAUAGCUGCGGAACUGGGCGC<br>CUGCUAUGCCAACAUAUUGCCAU | Processing |
| WT pre-miR-31-G45C | AGGCAAGAUGCUGGCAUAGCUGUUGAACUGGGAAC<br>CUCCUAUGCCAACAUAUUGCCAU | Processing |
| WT pre-miR-31-G45C/C46G | AGGCAAGAUGCUGGCAUAGCUGUUGAACUGGGAAC<br>CUCGUAUGCCAACAUAUUGCCAU | Processing |
| WT pre-miR-31-Δ43 | AGGCAAGAUGCUGGCAUAGCUGUUGAACUGGGAAC<br>UGCUAUGCCAACAUAUUGCCAU | Processing |
| WT pre-miR-31- Δ43/U44A | AGGCAAGAUGCUGGCAUAGCUGUUGAACUGGGAAC<br>AGCUAUGCCAACAUAUUGCCAU | Processing |
| WT pre-miR-31-U30C/A41G | AGGCAAGAUGCUGGCAUAGCUGCUGAACUGGGAGC<br>CUGCUAUGCCAACAUAUUGCCAU | Processing |
| WT pre-miR-31-G29A/C42U | AGGCAAGAUGCUGGCAUAGCUAUUGAACUGGGAAU<br>CUGCUAUGCCAACAUAUUGCCAU | Processing |

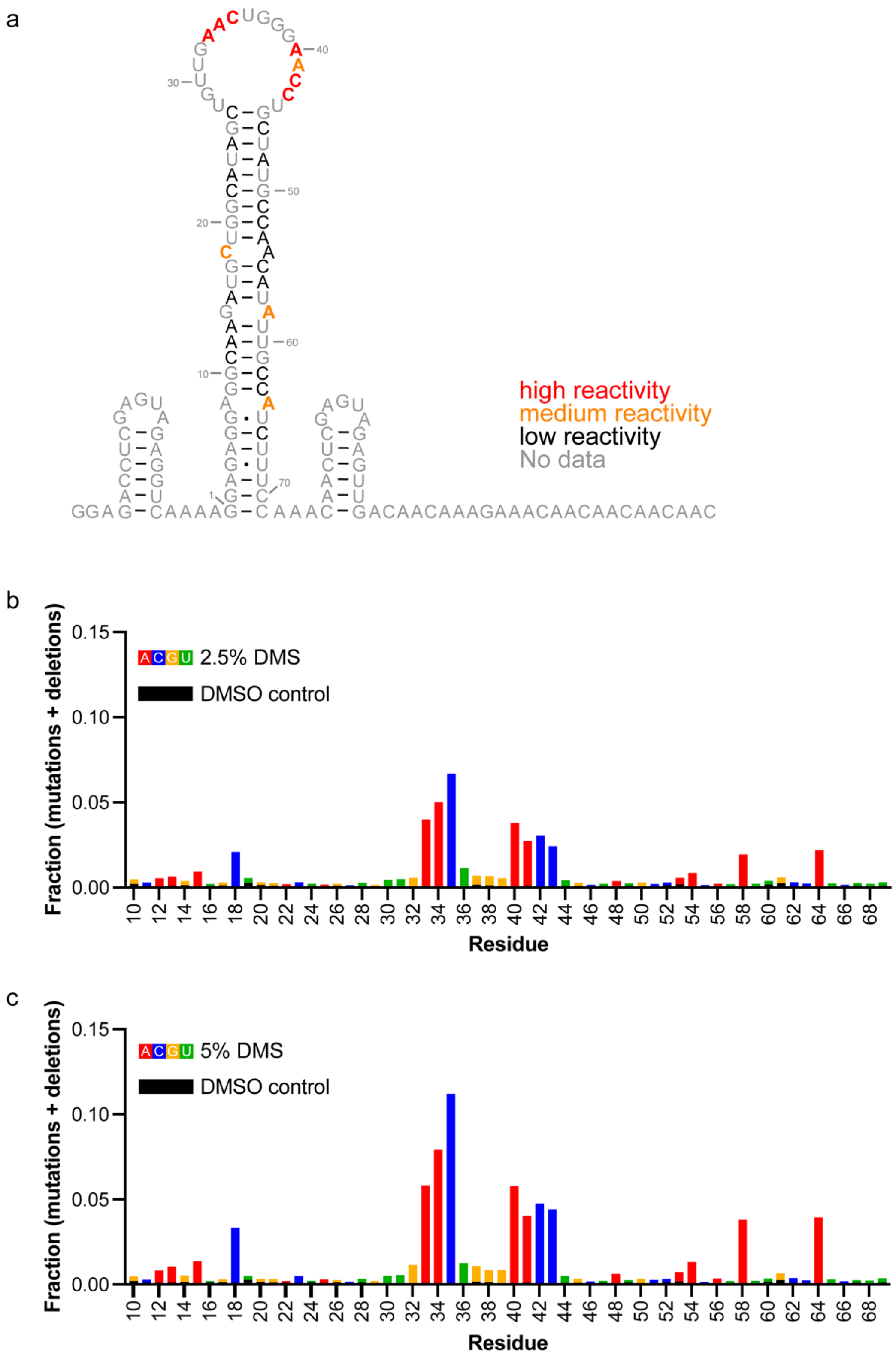


**Fig. S1. *In vitro* DMS-MaPseq of pre-miRNA-31 RNA. a)** Construct design and reactivity scores. 5′ and 3′ extensions were included to facilitate library preparation and sequencing. These regions were not predicted to disrupt the folding of pre-miR-31. **b** and **c)** Fraction of mutations and deletions on a per-residue basis upon reaction with 2.5% DMS (**b**) or 5% DMS (**c**). A control where DMSO was included in the reaction rather than DMS indicates minimal background mutations (black bars in **b** and **c**). Secondary structure was rendered using RNA2Drawer.[3]

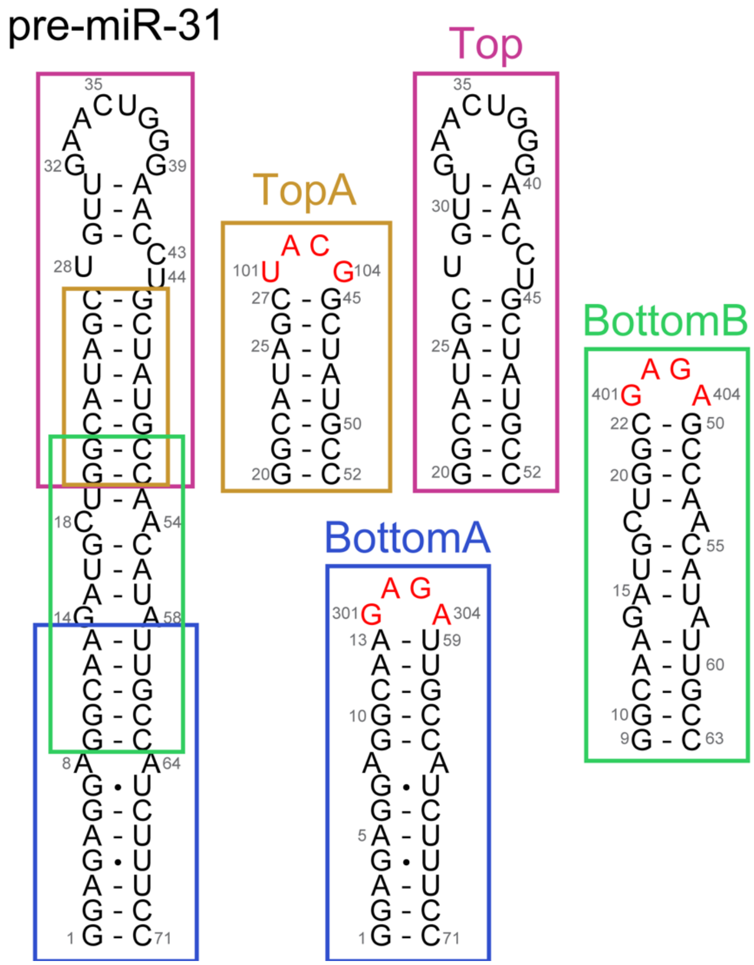


**Fig. S2. Oligo controls for NMR chemical shift assignment of FL-pre-miR-31.** Four oligonucleotide controls were designed to cover the entire FL-pre-miR-31 sequence. Non-native tetraloops (red) were included to cap oligos that truncated the apical loop. Secondary structures were rendered using RNA2Drawer.[3]

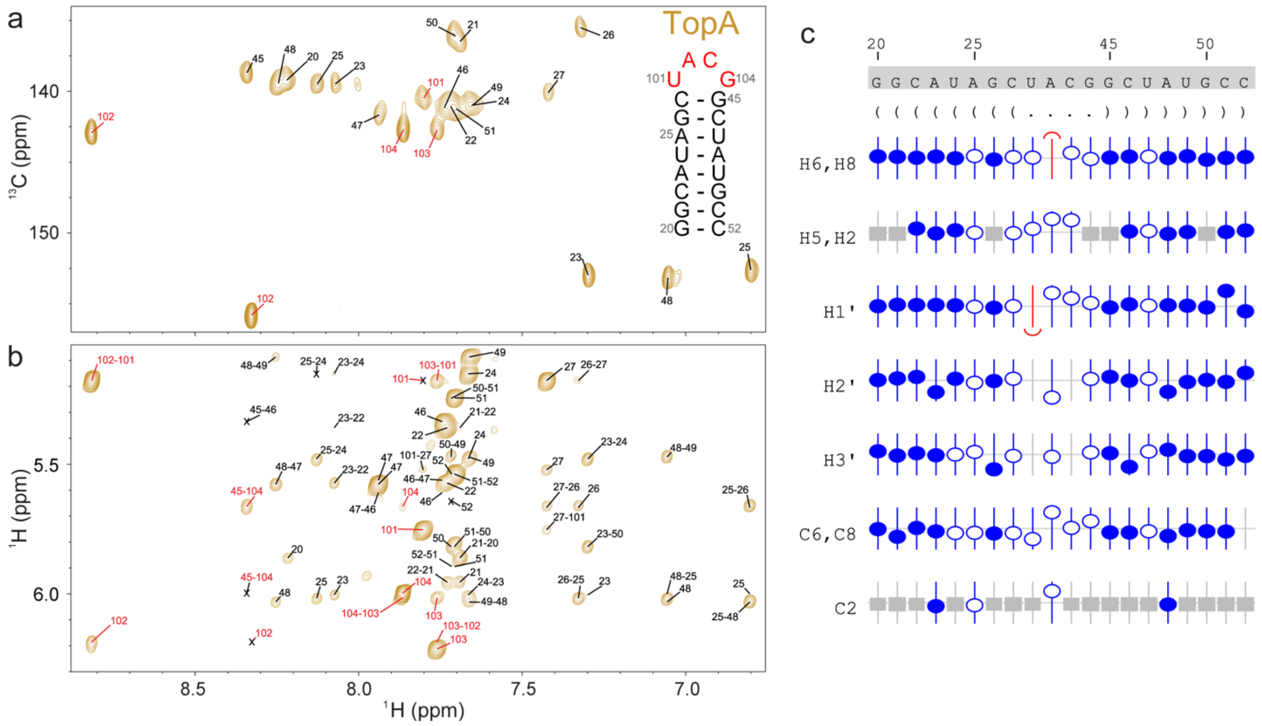


**Fig. S3. Assigned chemical shifts of TopA RNA. a)** $^{1}$H-$^{13}$C HMQC and **b)** $^{1}$H-$^{1}$H NOESY of TopA RNA. The signals assigned to GAGA tetraloop are colored red. NMR spectra were recorded at 0.4 mM RNA concentration, 50 mM K-phosphate buffer, pH=7.5, 1 mM $MgCl_2$ and 100% $D_2O$. **c)** Sequence analysis and validation of TopA chemical shift assignments. Nucleotide numbering, sequence, and secondary structure in Vienna format for TopA RNA. NMRViewJ chemical shift prediction software was used to validate proton (H6/H8, H5/H2, H1´, H2´, H3´) and carbon (C6/C8, C2) assignments. Assigned atoms are represented with blue circles (open and closed), while grey boxes denote atoms that are not present in a given base. Deviation from the predicted chemical shift is shown with the offset from the center. Filled circles indicate that there are chemical shifts for atoms with the same set of attributes in the BMRB. Open circles indicate atoms that have a prediction, but for which no exact matches of the attributes are available in the BMRB. Secondary structures were rendered using RNA2Drawer.[3]

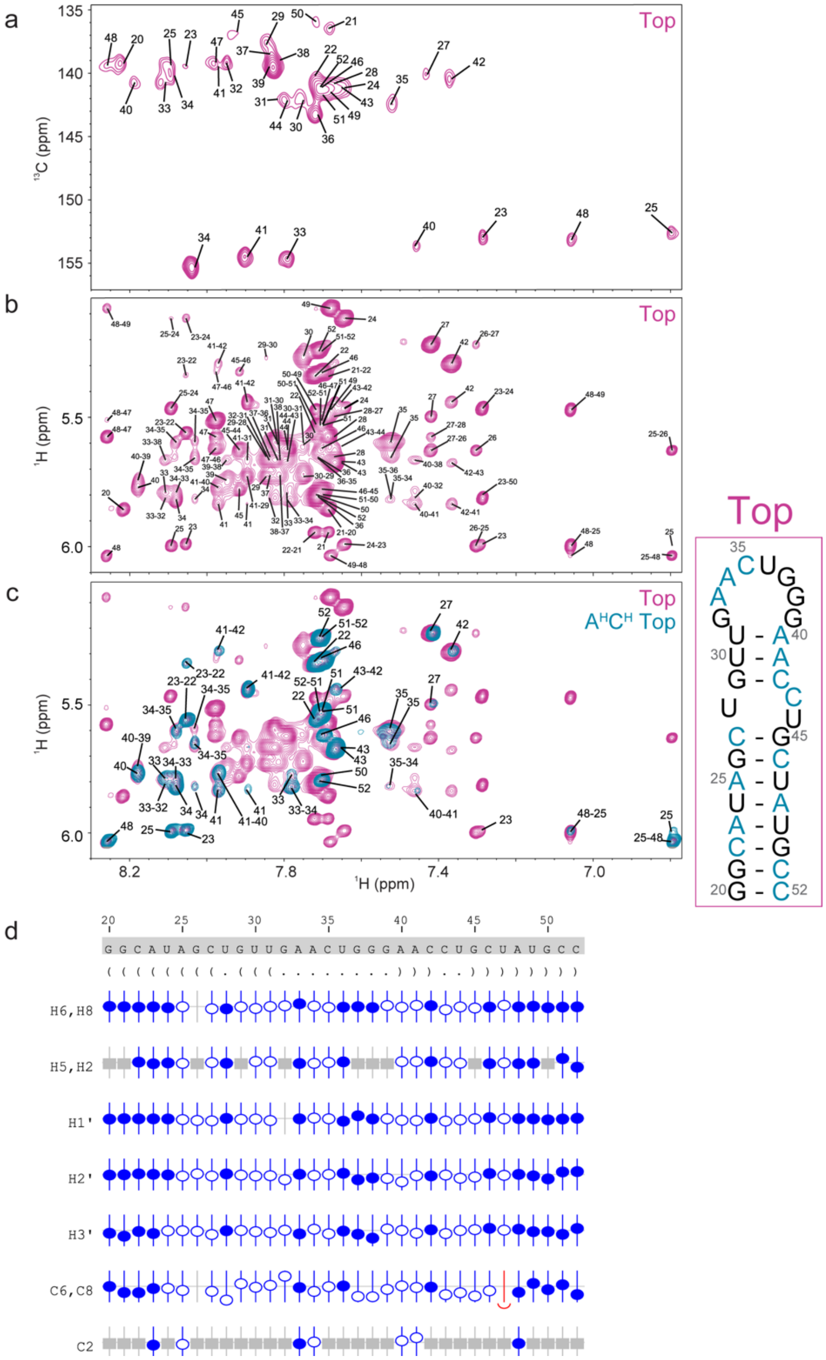


**Fig. S4. Assigned chemical shifts of Top RNA. a)** $^{1}$H-$^{13}$C HMQC and **b)** $^{1}$H-$^{1}$H NOESY of Top RNA. **c)** $^{1}$H-$^{1}$H NOESY spectrum overlay of fully-protiated (pink) and $A^{H}C^{H}$-labeled (teal) Top RNA. Secondary structure is colored to indicate the proton position (teal) in the $A^{H}C^{H}$-labeled Top RNA sample. All other sites are perdeuterated (black). NMR spectra were recorded at 0.4 mM RNA concentration, 50 mM K-phosphate buffer, pH=7.5, 1 mM $MgCl_2$ and 100% $D_2O$. **d)** Sequence analysis and validation of Top chemical shift assignments. Nucleotide numbering, sequence, and secondary structure in Vienna format for Top RNA. NMRViewJ chemical shift prediction software was used to validate proton (H6/H8, H5/H2, H1´, H2´, H3´) and carbon (C6/C8, C2) assignments. Assigned atoms are represented with blue circles (open and closed), while grey boxes denote atoms that are not present in a given base. Deviation from the predicted chemical shift is shown with the offset from the center. Filled circles indicate that there are chemical shifts for atoms with the same set of attributes in the BMRB. Open circles indicate atoms that have a prediction, but for which no exact matches of the attributes are available in the BMRB.

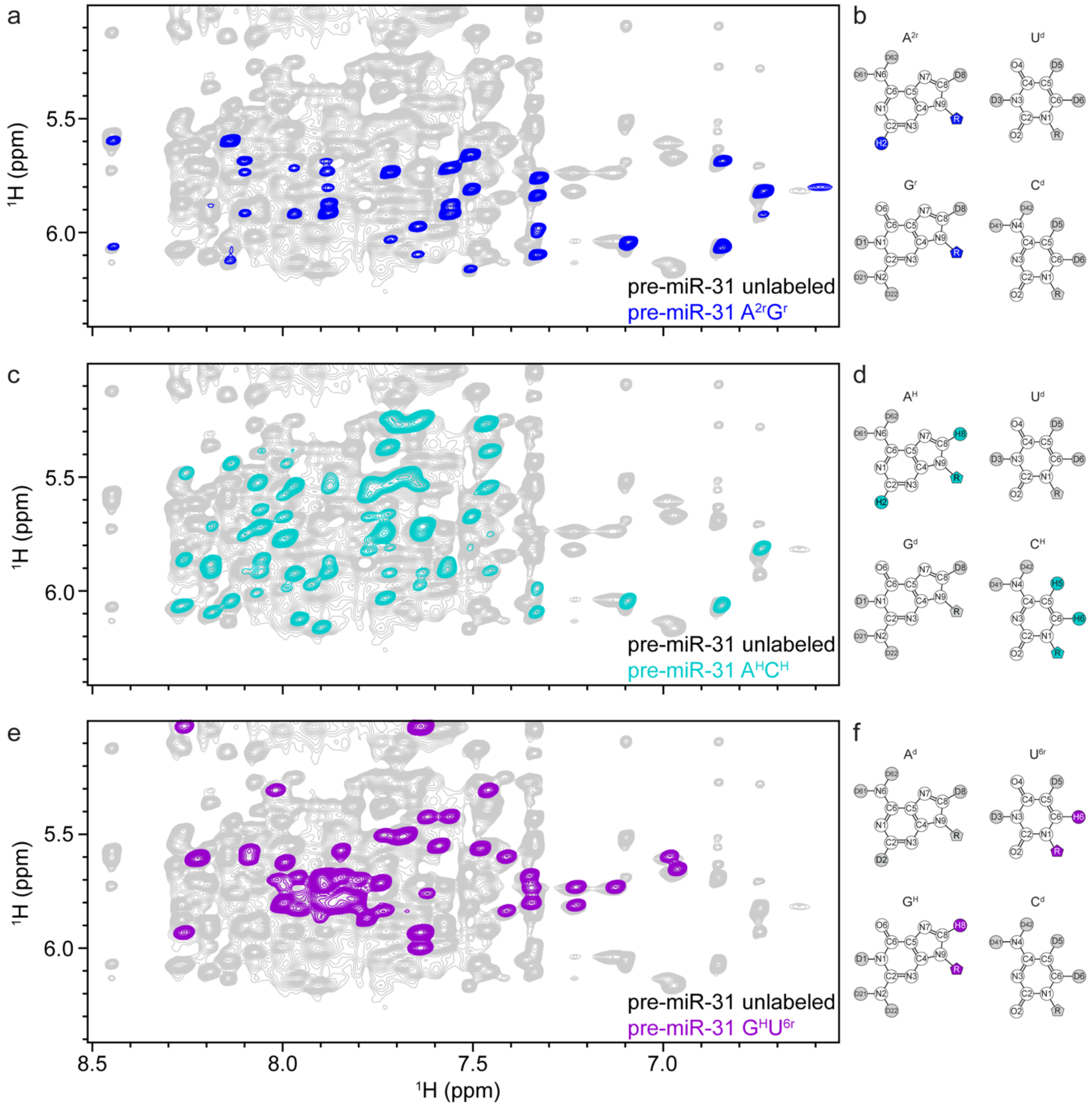


**Fig. S5. Deuterium labeling improves spectral quality by reducing overlap. a)** Overlay of the unlabeled (fully protiated, gray) and $A^{2r}G^{r}$-labeled (blue) pre-miR-31 $^{1}$H-$^{1}$H NOESY spectra. **b)** Chemical structures of the four nucleosides indicating sites of selective deuteration (gray shade). Sites containing non-exchangeable protons are colored blue. **c)** Overlay of the unlabeled (fully protiated, gray) and $A^{H}C^{H}$-labeled (teal) pre-miR-31 $^{1}$H-$^{1}$H NOESY spectra. **d)** Chemical structures of the four nucleosides indicating sites of selective deuteration (gray shade). Sites containing non-exchangeable protons are colored teal. **e)** Overlay of the unlabeled (fully protiated, gray) and $G^{H}U^{6r}$-labeled (purple) pre-miR-31 $^{1}$H-$^{1}$H NOESY spectra. **f)** Chemical structures of the four nucleosides indicating sites of selective deuteration (gray shade). Sites containing non-exchangeable protons are colored purple.

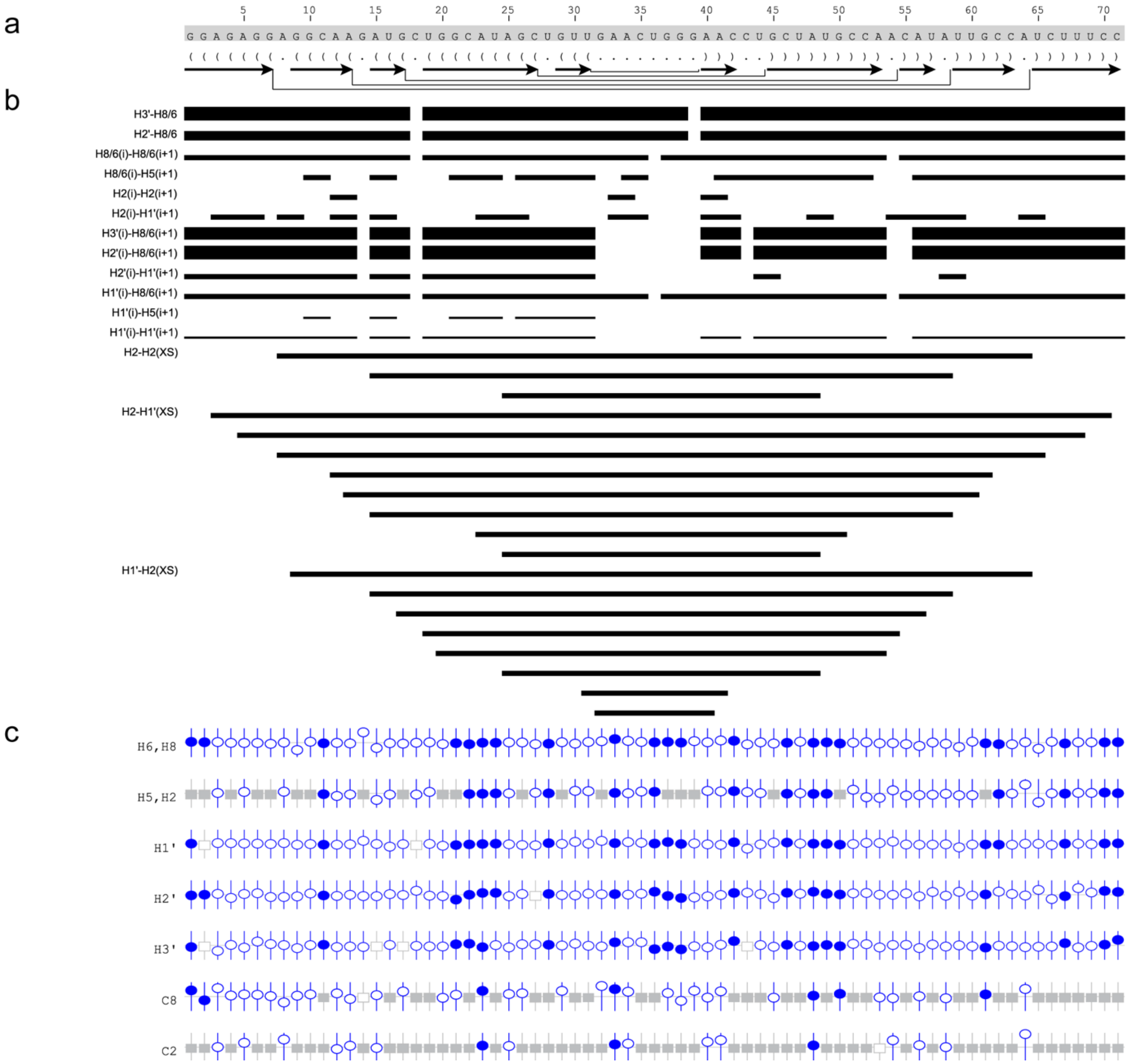


**Fig. S6. Summary of the secondary structure, NOE connectivity, and chemical shift assignment validation for FL pre-miR-31. a)** The secondary structure is shown beneath the sequence in Vienna format along with arrows to denote helical regions. **b)** NOE upper limit restraints for specified proton pairs used in CYANA and AMBER calculations are drawn as black bars. The thickness of the bar is representative of the strength of the measured NOE. **c)** NMRView chemical shift prediction software was used to validate assignments of H6/H8, H5/H2, H1′, H2′, H3′, C8 and C2. Protons that have been assigned in FL pre-miR-31 are indicated with blue circles (open and closed). Deviation from the predicted chemical shift is represented by deviation from the center. Predictions that are based on examples in the database of chemical shifts are shown as filled circles, predictions without data are shown as open circles. Filled grey squares are present in for nucleotides that do not contain a given proton or carbon. Unassigned resonances have open grey symbols.

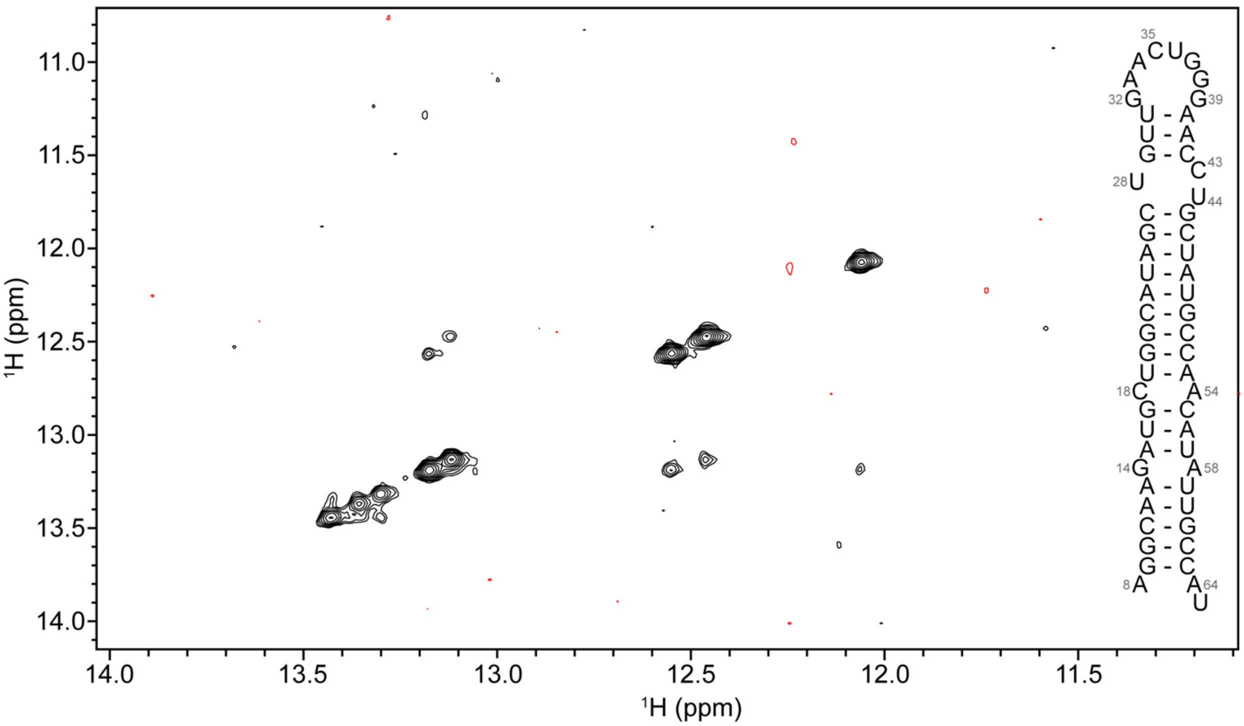


**Fig. S7. Imino proton NOESY spectrum of pre-miR-31.** The imino proton NOESY spectrum (mixing time 300 ms) of pre-miR-31 exhibited severe line broadening. The NMR spectrum was recorded at 0.33 mM RNA concentration, 50 mM K-phosphate buffer, pH=7.5, 1 mM $MgCl_2$ and 90% $H_2O$/10% $D_2O$ at 37°C and 600 MHz.

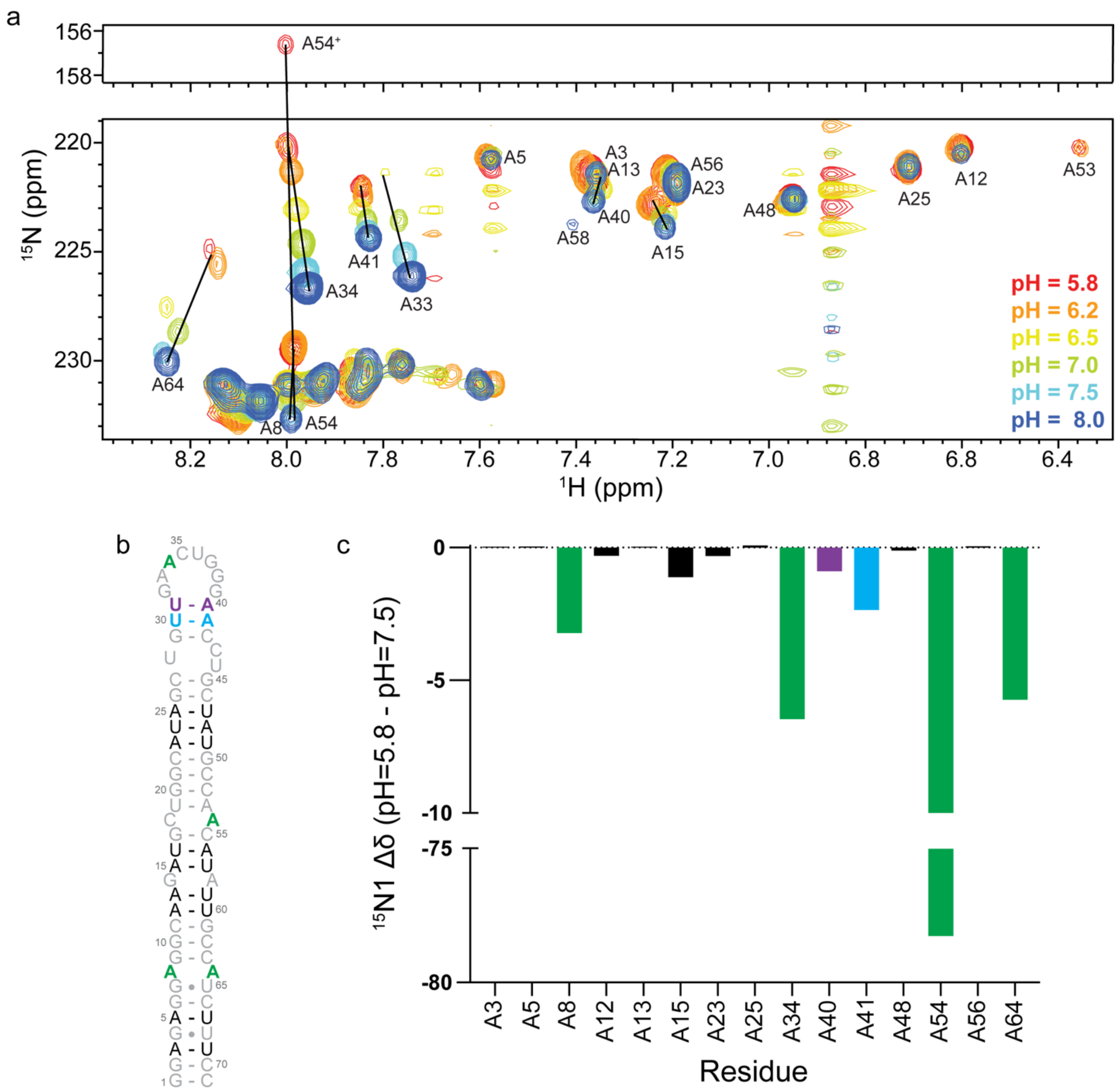


**Fig. S8. pH-dependence of unpaired adenosines. a)** BEST selective $^{1}$H-$^{15}$N HSQC spectra of $^{15}$N-AU labeled FL pre-miR-31, collected at various pH conditions. **b)** Secondary structure of FL pre-miR-31. **c)** Quantification of chemical shift perturbations (pH=5.8 – pH=7.5). A33 and A58 were not included in this analysis. The cross-peak of A33 is too broad to detect at pH=5.8 and cross-peak of A58 is severely overlapped at pH=5.8. Coloring follows the secondary structure in panel b.

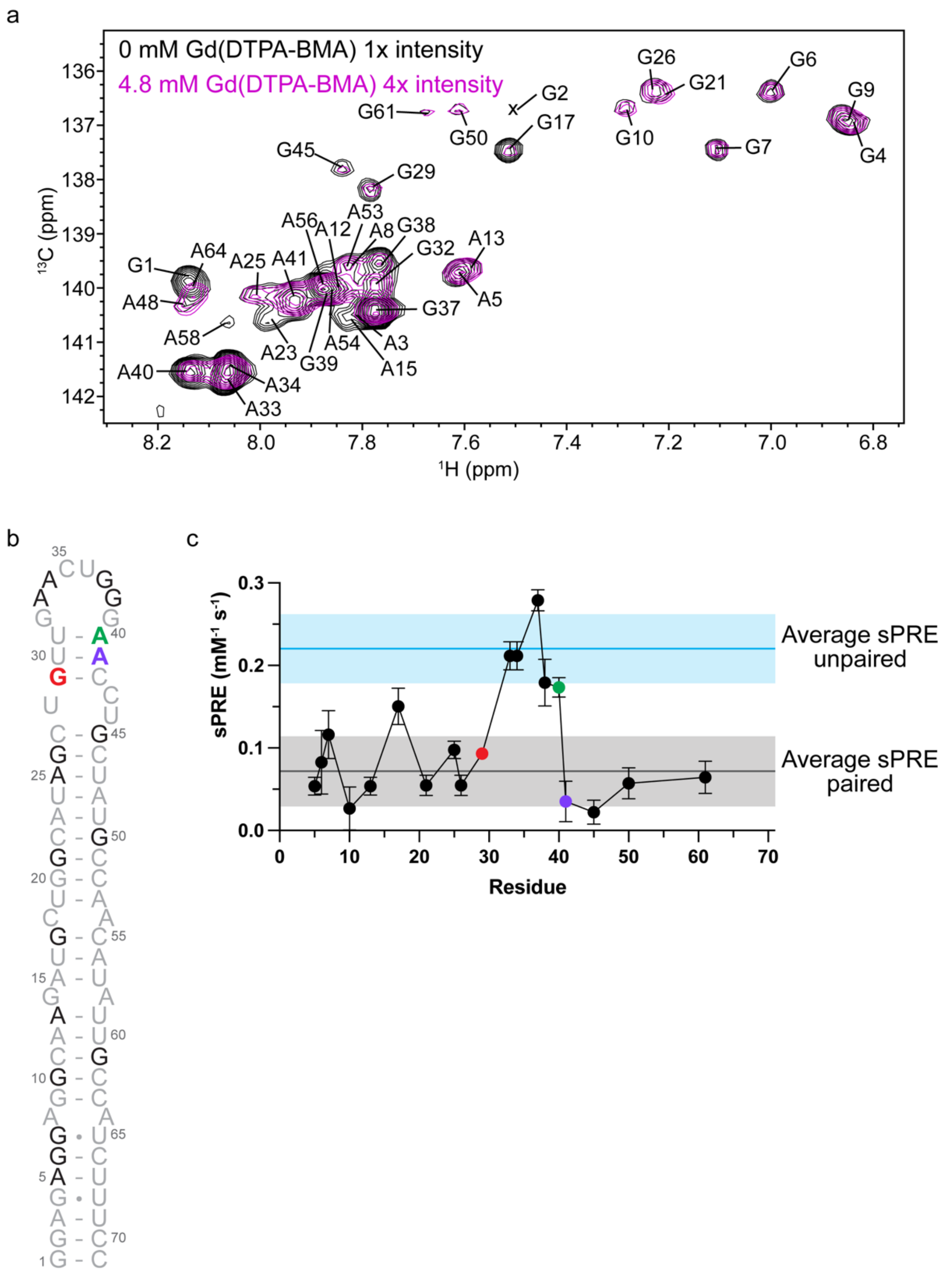


**Fig. S9. Solvent paramagnetic relaxation effect analysis of FL pre-miR-31 reveals solvent accessibility in the loop region.** a) $^{1}$H-$^{13}$C HSQC spectra of $^{15}$N/$^{13}$C A,G-labeled FL pre-miR-31 in the absence (black) and in the presence (magenta) of 4.8 mM paramagnetic compound Gd(DTPA-BMA). NMR spectra were recorded at 0.48 mM RNA concentration, 50 mM K-phosphate buffer (pD=7.5), 1 mM $MgCl_2$ and 100% $D_2O$ at 37 °C and 800 MHz. The H8-C8 assignments are labeled on the spectra. b) FL pre-miR-31 secondary structure colored to indicate the position of junction residues G29, A40 and A41 (red, green, and purple, respectively).

Residues shaded gray were not included in the analysis. c) sPRE data for aromatic H8 protons of FL pre-miR-31. The errors of the sPRE values were obtained from the linear regression as described previously.[4] The average sPRE values ± one standard deviation for unpaired (blue shading) and paired (grey shading) residues are indicated.

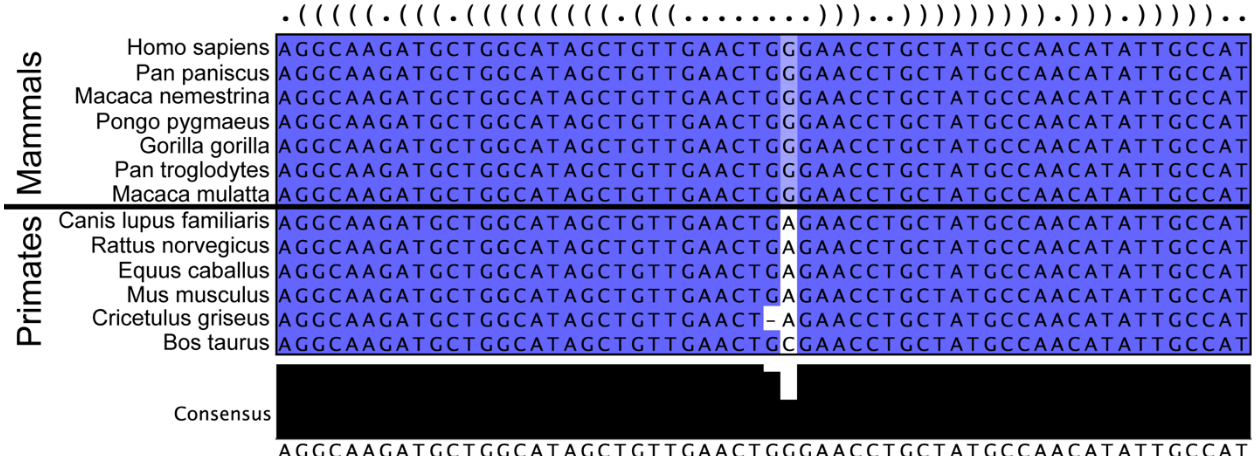


**Fig. S10. Sequence conservation of pre-miR-31.**

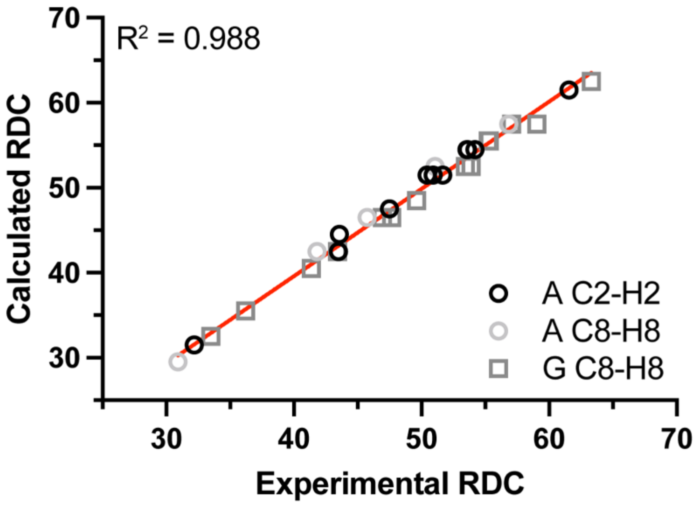


**Fig. S11. Correlation plot between measured and back-calculated RDCs for the lowest energy pre-miR-31 structure.**

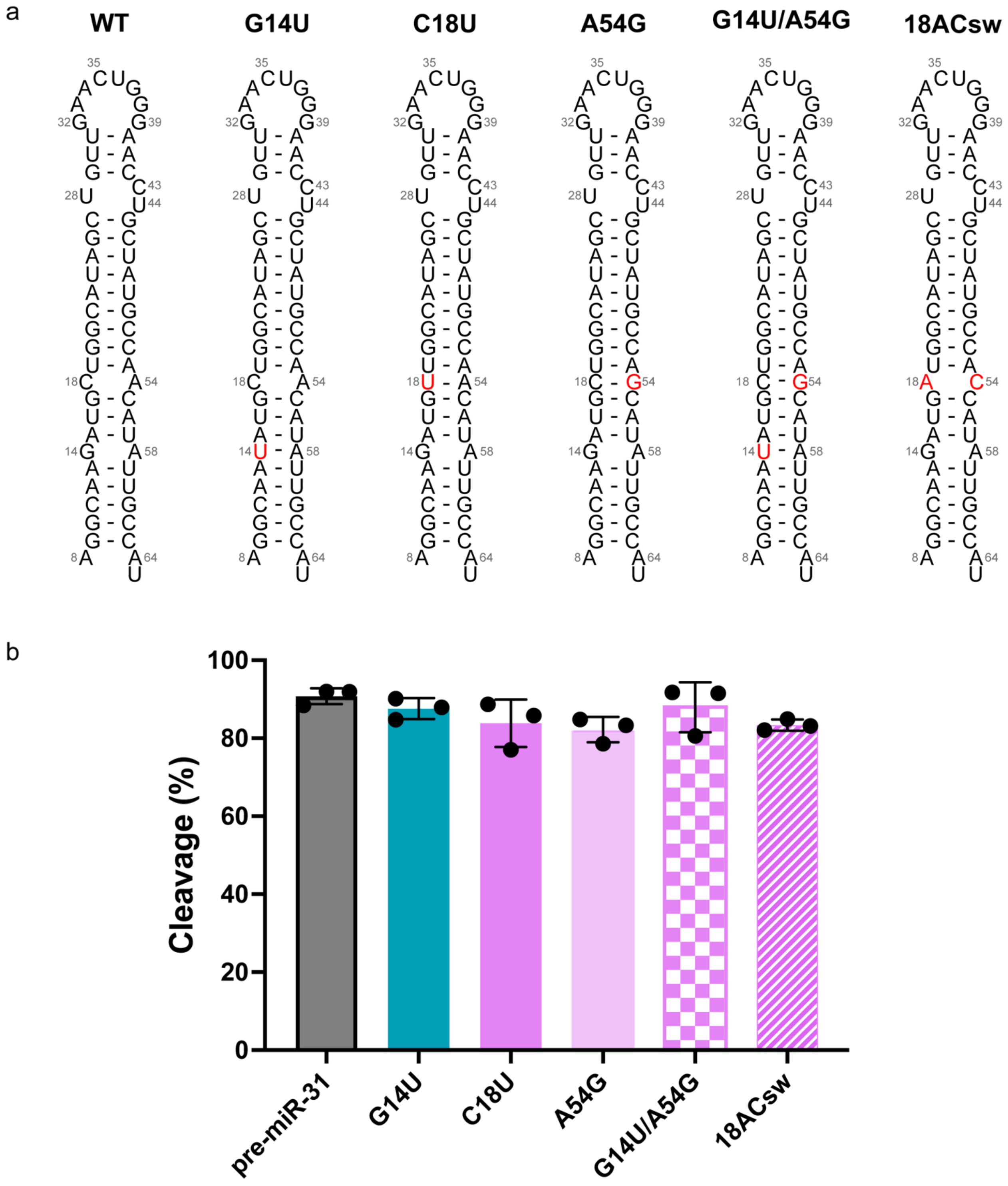


**Fig. S12. Mismatches in the stem region have no impact on Dicer processing. a)** Secondary structures of constructs designed to stabilize or destabilize the stem mismatches. Mutations are indicated with red lettering. **b)** Quantification of the Dicer processing efficiencies of pre-miR-31 RNAs at 10 min. Average and standard deviation from n=3 independent assays are presented. Individual replicates shown with black circles.

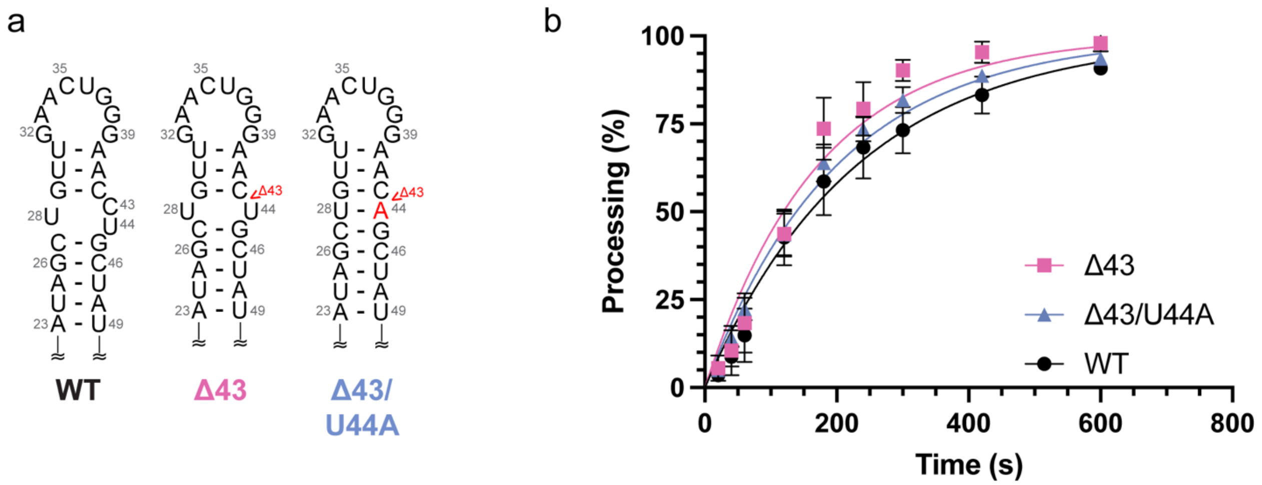


**Fig. S13. Mutations that minimize the internal loop at the dicing site are processed more efficiently than WT. a)** Secondary structures of constructs designed to minimize the internal loop at the dicing site. Mutations are indicated with red lettering. **b)** Quantification of the Dicer processing efficiencies of pre-miR-31 RNAs as a function of reaction time. Solid lines represent a best fit to a one-phase association equation. Values shown are the average and standard deviation from n=3 independent experiments.

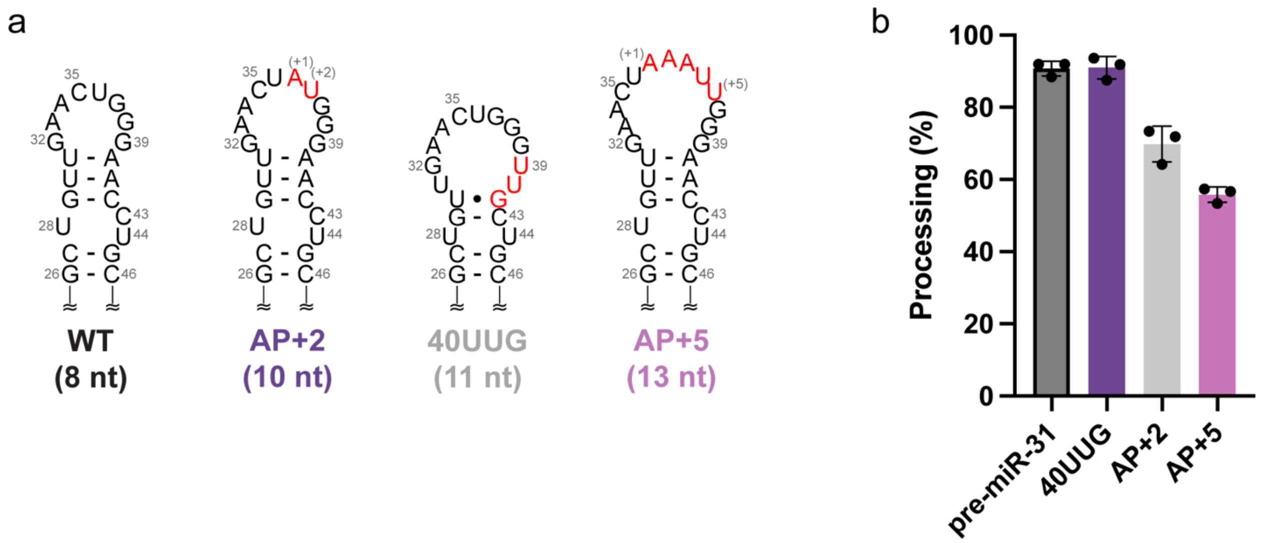


**Fig. S14. A two base-pair junction between the apical loop and dicing site recovers reduced processing efficiency due to large apical loop size. a)** Secondary structures of WT, AP+2, 40UUG, and AP+5 pre-miR-31 RNAs which have 8, 10, 11, and 13 nucleotide apical loops, respectively. Mutations are indicated with red lettering. **b)** Quantification of the Dicer processing efficiencies of pre-miR-31 RNAs at 10 min. Average and standard deviation from n=3 independent assays are presented. Individual replicates shown with black circles.

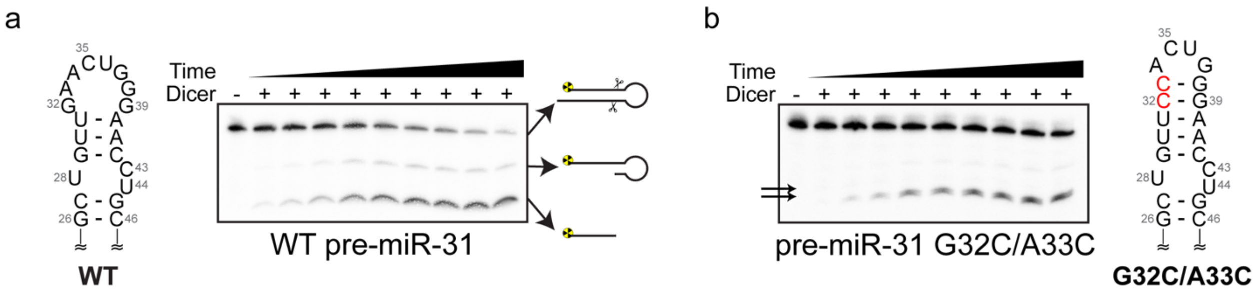


**Fig. S15. Extending the length of the helical junction between the apical loop and dicing site reduces processing efficiency and accuracy. a)** Secondary structure and Dicer processing gel of WT pre-miR-31. **b)** Secondary structure and Dicer processing gel of pre-miR-31 G32C/A33C. Two mature products are detected (black arrows).

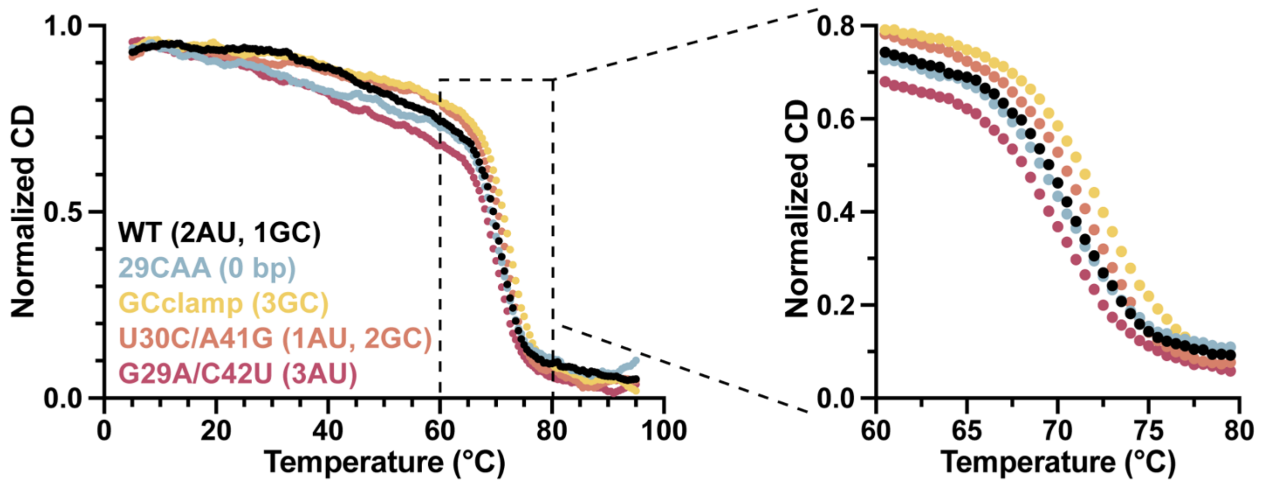


**Fig. S16. Thermal stability of junction mutants.** RNA thermal denaturation monitored by circular dichroism.
